# Acoustic Context Shapes Cortical Encoding of Spectral Discrimination in Normal and Impaired Hearing

**DOI:** 10.64898/2026.10.10.758219

**Authors:** Jaime A. Undurraga, Ioana Stefancu, Mats Rekswinkel, Lisbeth Birkelund Simonsen, Maaike Van Eeckhoutte

## Abstract

The auditory change complex (ACC) provides a cortical measure of acoustic discrimination, yet the mechanisms shaping responses to spectrotemporally complex sounds remain poorly understood. We investigated how acoustic context and temporal-envelope variability influence cortical encoding of spectral contrast in normal-hearing (NH) and hearing-impaired (HI) listeners. Electroencephalography (EEG) was used to record ACCs and auditory steady-state responses (ASSRs) evoked by spectrotemporally modulated stimuli. We compared conventional spectro-temporal modulations (STMs) with a novel discrete spectro-temporal modulation (DSTM) paradigm, examined temporal roving, and evaluated cortical responses to spectral contrast in NH and HI listeners. ACCs elicited by DSTM stimuli were larger and more symmetrical across transition directions than those elicited by conventional STMs. Temporal roving markedly reduced responses to intensity changes while largely preserving responses to spectral changes, indicating a partial dissociation between intensity-driven and spectrally driven cortical discrimination. ACC responses decreased systematically with spectral contrast and were weaker and delayed in HI listeners. By contrast, ASSRs remained highly detectable and showed no systematic dependence on spectral contrast, supporting a distinction between contrast-sensitive cortical discrimination and ongoing envelope encoding. Temporal roving further enhanced separation between NH and HI listeners. ACC response strength was significantly associated with behavioural STM discrimination thresholds and explained variance beyond age and audiometric thresholds. These findings demonstrate that cortical representations of spectral discrimination are shaped by acoustic context and temporal-envelope statistics. Temporal roving attenuates intensity-driven responses while preserving spectrally driven discrimination, providing insight into cortical encoding of spectrotemporal contrast and its relationship to perceptual performance.

## Introduction

Spectral and temporal acoustic cues are fundamental to auditory perception. They enable listeners to identify behaviourally relevant sounds, segregate competing sources, and organise information within complex acoustic environments, thereby supporting speech understanding, auditory scene analysis, and sound-source segregation (Bregman, 1990; Rabinowitz et al., 2013). Impaired representation of these cues is associated with degraded speech understanding, particularly in hearing-impaired (HI) listeners, highlighting the importance of spectrotemporal processing for everyday communication (Bernstein et al., 2013; Mehraei et al., 2014; Zaar et al., 2023). However, neural representations of auditory information are continuously shaped by the statistical context in which sounds occur, allowing the auditory system to maintain sensitivity across changing acoustic environments (Dean et al., 2005; Herrmann et al., 2013a; Espejo et al., 2019; Willmore and King, 2022). Understanding how such contextual information influences cortical representations of spectrotemporal contrast remains an important challenge in auditory neuroscience.

A variety of behavioural approaches have been developed to quantify sensitivity to spectrotemporal structure. Among these, spectro-temporal modulation (STM) detection paradigms have shown particular promise because they capture acoustic features that are prominent in natural speech. Behavioural sensitivity to STMs predicts speech-in-noise performance in HI listeners and provides information that complements conventional pure-tone audiometry (Zaar et al., 2023; Zaar et al., 2024a; Zaar, Simonsen, et al., 2024b). Similarly, discrimination of spectral-ripple stimuli has been associated with speech-perception abilities in both normal-hearing (NH) and HI listeners (Sohier et al., 2021). Collectively, these findings indicate that sensitivity to spectrotemporal structure constitutes an important dimension of auditory function that is not fully captured by threshold-based measures of hearing. Characterising its cortical representation is therefore important for understanding the neural mechanisms supporting auditory discrimination, with potential implications for objective assessment in clinical populations.

One approach for investigating cortical sensitivity to acoustic differences is the auditory change complex (ACC), a cortical evoked response elicited by a detectable change within an ongoing sound and measured non-invasively using electroencephalography (EEG) or magnetoencephalography (MEG). The ACC is typically characterised by a P1-N1-P2 complex occurring approximately 50, 100, and 200 ms after change onset, respectively (Martin and Boothroyd, 2000). ACCs can be elicited by changes in frequency, spectral content, temporal structure, and intensity (Dimitrijevic et al., 2009; Dimitrijevic et al., 2011; Nishihara et al., 2011; Han and Dimitrijevic, 2015; Vonck et al., 2019; Calcus et al., 2022; Fujita et al., 2022; Wang et al., 2025). They have also been used to investigate discrimination of more complex acoustic features, including vowels, spectral ripples, and spectrotemporal modulations (Martin and Boothroyd, 2000; Martin et al., 2010; Won et al., 2011; Cheek and Cone, 2020; Sohier et al., 2021; Jabeen et al., 2025; Simonsen et al., 2025; Simonsen et al., 2026; Rekswinkel et al., 2026).

Because the response provides an objective measure of cortical sensitivity to acoustic changes, the ACC has become an attractive tool for investigating neural mechanisms of auditory discrimination. Moreover, associations between ACC measures and perceptual performance suggest that ACCs provide insight into the cortical processes supporting auditory perception (see Meehan et al., 2025, for a review).

However, the ACC primarily reflects the neural response elicited by a discrete change event and does not directly characterise the ongoing representation of the stimulus. Auditory steady-state responses (ASSRs) provide a complementary measure of phase-locked neural encoding of periodic stimulus-envelope modulations (Picton et al., 2003). Recording ACCs and ASSRs simultaneously therefore allows change-related cortical discrimination to be evaluated alongside ongoing envelope encoding. This comparison helps determine whether contextual effects observed in the ACC are accompanied by corresponding changes in the neural representation of the stimulus envelope.

Nevertheless, the factors governing ACC generation remain incompletely understood, particularly when spectrotemporally complex sounds are used.

ACC responses are often interpreted as reflecting sensitivity to the stimulus feature being experimentally manipulated. However, changes in spectrotemporal structure can alter several stimulus dimensions simultaneously. For example, spectrotemporal manipulations may change perceived loudness in addition to the statistical properties of the acoustic context. Consequently, the measured cortical response may reflect a combination of spectral discrimination, intensity-related processing, and adaptive neural mechanisms that depend on recent stimulation history. Distinguishing among these contributions is essential for determining what information ACC responses encode and how reliably they index cortical sensitivity to a particular acoustic feature.

Recent work by Simonsen et al. (2025) demonstrated that ACCs elicited by spectrotemporally modulated sounds depend strongly on both the spectral characteristics of the stimulus conveying the modulations—carrier—and the direction of the acoustic transition. Responses were substantially larger when the modulations were conveyed by a tonal carrier than by a noise carrier. They were also larger for transitions from an unmodulated reference to a spectrotemporally modulated target than for the reverse transition. These findings indicate that cortical discrimination responses are not determined solely by the physical magnitude of an acoustic change. Instead, the neural representation of a change appears to depend on the spectrotemporal context in which it occurs.

One possible mechanism underlying these context effects is adaptive coding. Auditory neurons continuously adjust their responsiveness according to the statistical properties of recent sensory inputs, including both the mean and variance of acoustic stimulation (Dean et al., 2005; Kvale and Schreiner, 2004; Nagel and Doupe, 2006; Dean et al., 2008; Wen et al., 2009; Wen et al., 2012; Bakay et al., 2018; Espejo et al., 2019; Willmore and King, 2022). Such adaptation is thought to optimise neural sensitivity to the prevailing sensory environment by adjusting the dynamic range over which changes are represented. Comparable phenomena have been observed in human auditory cortex, where neural responses depend on the statistical distribution of preceding sounds (Herrmann et al., 2013a; Herrmann et al., 2013b; Herrmann et al., 2014). Changes in contextual variability may therefore modify the neural representation against which subsequent acoustic events are evaluated, altering the magnitude or timing of the resulting cortical discrimination response.

A second, non-exclusive mechanism is response-rate adaptation. Repeated or sustained neural activation can reduce responsiveness to temporally adjacent events as a result of refractoriness, synaptic fatigue, or the persistence of an auditory sensory-memory trace (Sams et al., 1993; Mcevoy et al., 1997; Rosburg, 2004; Herrmann et al., 2016; Schellekens et al., 2025). Increasing the interval between successive acoustic transitions would be expected to reduce these effects and enhance the corresponding cortical responses. Together, context-dependent coding and response-rate adaptation predict that the cortical representation of a spectral change should depend not only on the physical properties of the change, but also on the statistical and temporal structure of the preceding acoustic context.

A related challenge is that cortical discrimination responses may contain substantial contributions from intensity-based processing. Relatively small changes in sound level are sufficient to elicit robust ACC responses (Martin and Boothroyd, 2000; Dimitrijevic et al., 2011; Nishihara et al., 2011; Fujita et al., 2022). ACCs evoked by spectrotemporal manipulations may therefore partly reflect intensity- or loudness-related cues rather than cortical representations of spectral differences per se. Attenuating these potential intensity-related contributions whilst preserving cortical responses driven by spectral information would provide a more specific measure of spectral discrimination.

One potential approach is to manipulate the temporal statistics of the stimulus. Natural sounds, particularly speech, exhibit pronounced low-frequency envelope fluctuations. Introducing similar temporal-envelope variability increases fluctuations in instantaneous intensity without altering the underlying spectral contrast. This variability may reduce the reliability of intensity-based contextual representations by preventing the formation of a stable reference level, whilst preserving spectral information relevant to discrimination. Temporal-envelope variability may therefore help dissociate intensity-driven and spectrally driven contributions to cortical change responses while simultaneously revealing how the statistics of the ongoing acoustic context shape cortical representations of spectral contrast.

The present study investigated how acoustic context, temporal-envelope variability, and hearing loss influence cortical responses to spectrotemporal contrast measured using EEG. First, we examined whether previously reported carrier- and direction-dependent ACC effects could be explained by context-dependent or response-rate-dependent mechanisms. We compared conventional STMs, in which spectral and temporal structure vary within the target segment, with a novel discrete spectro-temporal modulation (DSTM) paradigm, in which the acoustic change is restricted to spectral structure, whilst manipulating carrier type, transition direction, and presentation rate. Second, we tested whether speech-like temporal-envelope variability differentially affects intensity-driven and spectrally driven contributions to cortical discrimination. Third, we characterised cortical sensitivity to spectral contrast in NH and HI listeners and examined how temporal-envelope variability shapes these representations. ASSRs were analysed in parallel to determine whether changes in ACC responses were accompanied by differences in ongoing envelope encoding and to evaluate potential effects of presentation level and stimulus audibility.

Finally, we evaluated the relationship between cortical response measures and behavioural discrimination performance assessed using the Audible Contrast Threshold (ACT) test (Zaar, Simonsen, et al., 2024b).

We hypothesised that cortical discrimination responses would depend strongly on the statistical properties of the ongoing and preceding acoustic context, consistent with context-dependent adaptive coding. We further hypothesised that speech-like temporal-envelope variability would selectively attenuate intensity-driven responses whilst preserving responses elicited by spectral contrast. By reducing competing intensity-related contributions, we expected temporal-envelope variability to produce a more systematic cortical representation of spectral contrast and strengthen the relationship between cortical and behavioural discrimination. Finally, we hypothesised that hearing impairment would be associated with weaker, delayed, and less differentiated cortical representations of spectral contrast.

## Methods

### 2.1. Participants

Forty-six participants were recruited for this study including 18 NH (20-44 years, 6 female) and 28 HI (60-81 years, 17 female) listeners. Participants were recruited through the Technical University of Denmark (DTU) and Rigshospitalet. Experimental procedures were approved by the Science-Ethics Committee for the Capital Region of Denmark (H-16036391), and written informed consent was obtained from all participants prior to participation. Participants received financial compensation for their time.

All NH participants demonstrated pure-tone hearing thresholds ≤ 20 dB hearing level (HL) between 250 Hz and 2 kHz, corresponding to the bandwidth of the experimental stimuli. HI participants had hearing thresholds of no more than 80 dB HL within the same frequency range (Figure 1a). Better-ear PTA was defined as the lower of the left- and right-ear pure-tone averages across 500, 1000, 2000, and 4000 Hz.

**Figure 1.**
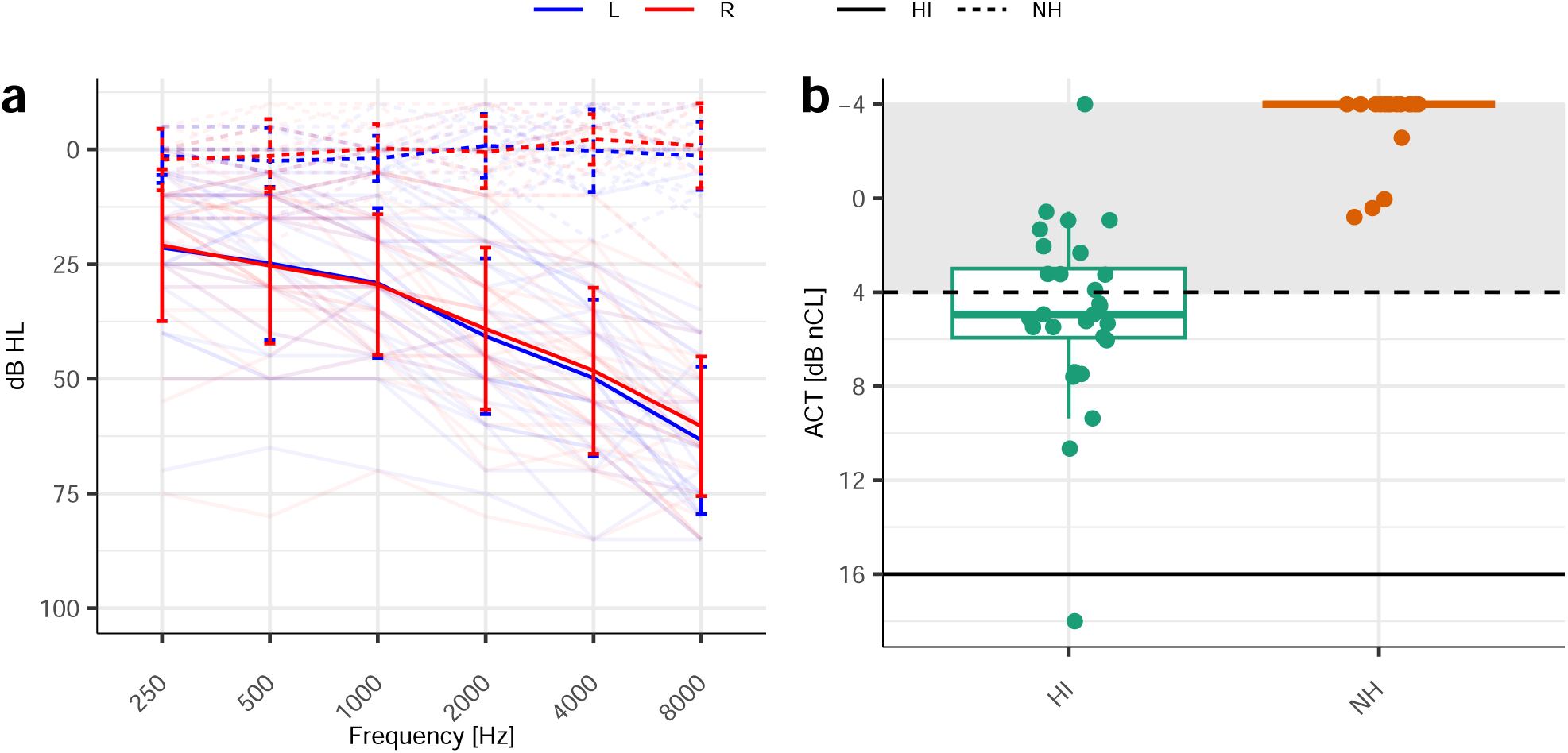
**a**, Pure-tone audiometric thresholds for NH (dashed lines) and HI (continuous lines) listeners for the left (blue) and right (red) ears, respectively. **b**, ACT values for NH and HI participants. The shaded area shows the range of normal performance (Zaar, Simonsen, et al., 2024b).

Pure-tone audiometry was conducted in a sound-attenuating booth using a calibrated clinical audiometer (Affinity Compact, Interacoustics, Denmark). Thresholds were obtained using the Hughson-Westlake ascending procedure with 5-dB step sizes. Participants were excluded if they reported neurological disorders or other conditions potentially affecting auditory processing, or if otoscopy revealed excessive cerumen.

### 2.2. Behavioural Assessment

All participants completed the ACT test (Affinity Compact, Interacoustics, Denmark), a behavioural measure of sensitivity to spectrotemporal modulations similar to those present in speech signals (Zaar et al., 2023; Zaar et al., 2024a; Zaar, Simonsen, et al., 2024b). The stimuli consisted of 1-s pink-noise bursts spanning 354-2000 Hz. Target stimuli contained combined spectral and temporal modulations (2 cycles/octave and 4 Hz modulation rate), producing a spectrotemporally modulated percept often described as “siren-like”. Reference stimuli had identical bandwidth and duration but contained no imposed modulation. Participants were instructed to identify the modulated stimulus by a button press. The modulation depth, expressed as normalised contrast level (nCL), in dB, was varied adaptively using a 2-down 1-up procedure with 2-dB step sizes until the Hughson-Westlake stopping criterion was met.

An ACT of 0 dB nCL corresponds to typical performance observed in young NH listeners, whereas 16 dB nCL represents the maximum modulation depth and therefore the poorest performance. To minimise the influence of audibility limitations, ear-specific amplification was applied according to individual audiometric thresholds following the procedures described by Zaar and colleagues (Zaar et al., 2023; Zaar et al., 2024a; Zaar, Simonsen, et al., 2024b).

### 2.3. Stimuli

#### 2.3.1. Carrier signals

Physiological responses were evoked using either a noise carrier (NC) or a tonal carrier (TC), following previous studies of cortical responses to spectrotemporal modulations (Simonsen et al., 2025). The NC consisted of 2500 sinusoidal components with equal amplitude, random initial phases, and frequencies distributed logarithmically across the stimulus bandwidth (Figure 2a,b,e,f). The TC consisted of 17 sinusoidal components with random initial phases and frequencies distributed uniformly on a linear frequency scale (Figure 2c,d,g-j). The spectral power density of the TC followed a 1/*f* distribution. All stimuli occupied a bandwidth of 337.1-2105.3 Hz. Spectrotemporal modulations were restricted to the range 354-2000 Hz, whilst frequency components outside this region remained unmodulated to minimise potential edge cues.

**Figure 2.**
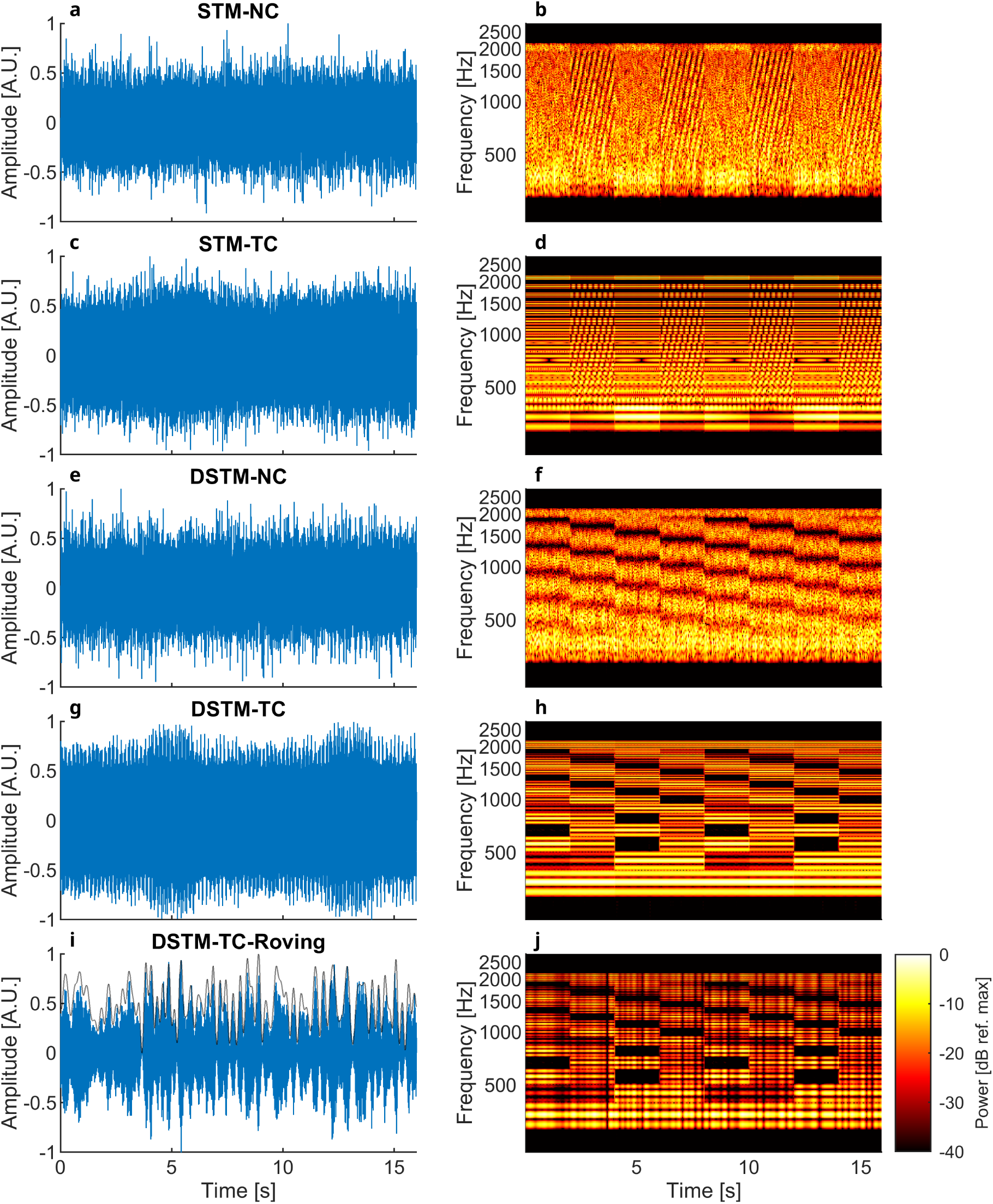
Time waveforms and spectrograms at 0 dB CL showing the power spectral density (PSD) for STM using a NC (**a&b**), STM using a TC (**c&d**), DSTM-90°using a NC (**e&f**), DSTM-90°using a TC (**g&h**), DSTM-90°using a TC and speech-like temporal roving (**i&j**; the modulator is shown by the solid black line). The transitions evoking the ACC—visible in the spectrograms—were imposed every 2 s in this example.

#### 2.3.2. Spectrotemporal Modulations

Two classes of stimuli were used throughout the study: continuously spectrotemporally modulated stimuli (STM) and discrete spectrotemporal modulation (DSTM) stimuli (Figure 2).

For STM stimuli, the signal alternated between a reference segment containing no spectrotemporal modulation and a target segment containing simultaneous spectral and temporal modulations (2 cycles/octave and 4 Hz, respectively). Thus, both spectral and temporal structure varied continuously within the target segment.

The DSTM stimulus was developed to evoke cortical responses to abrupt spectral transitions whilst maintaining a constant spectrotemporal structure within each segment. DSTM segments consisted of static spectral ripples (2 cycles/octave). Consecutive segments differed by 90° phase shift in the spectral ripple pattern (DSTM-90), thereby producing an instantaneous change in spectral content at the segment boundary. This design generated abrupt spectral transitions similar to those employed by Sohier et al. (180°), whilst preserving identical average spectral energy across segments. If not explicitly stated, the level of spectro-temporal modulation, here referred to as contrast level (CL), was 0 dB CL, corresponding to the maximum modulation depth.

Transition periods of 2.0 or 3.3 s were employed depending on the experiment.

#### 2.3.3. Temporal Roving

To manipulate temporal-envelope variability, a subset of DSTM stimuli was additionally modulated by a low-frequency, speech-like amplitude envelope (0.2-4 Hz), referred to as temporal roving (Figure 2i&j). This modulation was applied continuously across the entire stimulus and introduced slow fluctuations in instantaneous sound level resembling those observed in natural speech (Ahissar et al., 2001).

Temporal roving was employed in Experiments 2 and 3 to examine how increased temporal-envelope variability affected cortical discrimination responses.

#### 2.3.4. Auditory Steady-State Response Modulation

To permit simultaneous recording of ASSRs, all stimuli were amplitude modulated at 42 Hz (Picton et al., 2003). Stimulus transitions were aligned with minima of the 42-Hz modulation envelope to minimise spectral splatter and other transient cues unrelated to the intended experimental manipulation. This manipulation resulted in additional side bands around each tonal carrier (Figure 2).

### 2.4. Audibility compensation

To ensure adequate audibility for HI participants, frequency-specific amplification was applied individually to each ear. The procedure followed the sufficient-audibility approach used in the ACT test (Humes, 2007; Zaar et al., 2023; Zaar et al., 2024a), with the objective of maintaining stimulus levels at least 15 dB above hearing threshold across the stimulus bandwidth.

Audiometric thresholds were interpolated onto one-third-octave band centre frequencies and combined with minimum audible field (MAF) values for diffuse-field listening (ISO 389-7, 2019). The spectral power density of each stimulus was estimated using Welch’s method and compared with the participant-specific audibility curve. Frequency-dependent gain was then applied where necessary to ensure supra-threshold audibility whilst limiting the level within each one-third-octave band to a maximum of 104 dB sound pressure level (SPL) for safety. The resulting gain function was implemented as a zero-phase finite impulse response (FIR) filter and applied individually to each participant and ear.

This procedure ensured comparable audibility of the stimuli across listeners whilst minimising the influence of peripheral hearing sensitivity on cortical response measurements.

### 2.5. EEG recordings

EEG recordings were conducted inside an electromagnetically and acoustically shielded booth. Stimuli and trigger signals were generated using the open-source MATLAB-based software “AEP-GUI” and delivered through a digital sound card (RME Fireface UC). The audio output was amplified (Phonitor Mini, SPL Audio, Germany) and presented through insert earphones (ER3A, Etymotic Research, USA).

Stimuli were calibrated to 65 dB SPL using a 2-cc coupler (G.R.A.S., Denmark) when neither audibility compensation nor temporal roving was applied. Following audibility compensation, overall presentation levels varied across HI participants. Application of temporal roving resulted in an average reduction of approximately 5 dB in overall stimulus level relative to the non-roving condition.

EEG recordings were acquired using a BioSemi ActiveTwo system (BioSemi, Amsterdam, The Netherlands) with 64 scalp electrodes positioned according to the international 10-20 system and two additional mastoid electrodes (TP9 and TP10). Data were sampled at 2048 Hz with 24-bit resolution. The BioSemi anti-aliasing filter provided a low-pass cutoff at 512 Hz. Electrode offsets were maintained below 40 mV throughout recording sessions.

A total of 160 stimulus epochs were collected per condition, corresponding to approximately 5.3 min of recording for conditions employing a 2-s transition interval.

### 2.6. EEG processing

All EEG analyses were performed using the open-source Python package “pEEGy” (Undurraga, 2026).

#### 2.6.1. EEG Preprocessing

Continuous EEG recordings were first re-referenced to Cz and polarity-inverted for visualization purposes. Data were subsequently downsampled to 500 Hz. Channels exhibiting large and persistent DC offsets were automatically identified and excluded from subsequent analyses. Typically, no more than one or two channels were removed per participant.

For ACC analyses, EEG signals were band-pass filtered between 3 and 20 Hz using an FIR Kaiser filter with 60 dB attenuation in the stopband and a 1-Hz transition bandwidth. For ASSR analyses, recordings were filtered between 2 and 200 Hz using the same filter design.

Following filtering, recordings were segmented into epochs aligned to the stimulus transitions.

To compare the effect of context preceding a change, analyses were carried out for epochs grouped into three conditions: reference-to-target (RT) transitions, target-to-reference (TR) transitions, or the combination of both transition directions (R&T). Note that we employ the terms ‘reference’ and ‘target’ as in the behavioural task and as originally employed in Simonsen et al. (2025). However, during EEG recordings participants were not actively detecting the presence of sound change. Instead, they watched a subtitled film of their choice whilst remaining as relaxed and still as possible. Importantly, RT and TR do not represent equivalent acoustic transitions in the STM and DSTM paradigms. In the STM stimulus, RT and TR transitions differ in the preceding spectrotemporal context because one segment contains ongoing spectrotemporal modulation whereas the other does not. In contrast, DSTM segments differ only in spectral-ripple phase, such that RT and TR labels primarily indicate transition direction rather than distinct acoustic contexts. Consequently, RT and TR have a more arbitrary interpretation in the DSTM analyses.

#### 2.6.2. ACC Analysis

ACC responses were quantified using the Fmpi method (Undurraga et al., 2026), which computes a response-strength metric (F-value), the probability that a response is present, and an estimate of post-average residual noise. The F-value was used as the primary measure of response detectability and can be interpreted as a signal-to-noise ratio (SNR) metric according to:

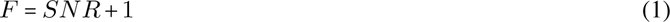

such that F-values closer to 1 indicate little or no evoked response.

ACC analyses were performed within a post-transition response window extending from 50 to 400 ms. Response amplitude was defined as the peak-to-peak difference between the N1 and P2 components. The N1 corresponded to the most prominent negative peak occurring within the response window, and P2 was defined as the largest positive peak occurring between the N1 latency and the end of the analysis window.

Response detection rates were determined using the statistical significance criterion provided by the Fmpi method, using Bonferroni correction when left and right mastoid channels were evaluated simultaneously.

#### 2.6.3. ASSR Analysis

ASSRs were elicited by the continuous 42-Hz amplitude modulation present in all stimuli. For each epoch, the spectral amplitude at 42 Hz was estimated using the fast Fourier transform (FFT). Residual noise was estimated from neighbouring frequency bins within ±2 Hz of the modulation frequency whilst excluding the target frequency bin.

ASSR response strength was quantified using the Fmpi method applied to the frequency-domain data. ASSR amplitude corresponded to the magnitude of the 42-Hz frequency component. ASSR latency was estimated from the phase of the 42-Hz response following established procedures for steady-state responses (Herdman et al., 2002; Undurraga et al., 2016). ASSRs were analysed separately for reference (R) segments, target (T) segments, and their combination R&T. For comparison with the ACC analyses and figures, R and T segment estimates were associated with the corresponding TR and RT recording blocks, respectively.

### 2.7. Experimental Design

#### 2.7.1. Experiment 1: Effects of Acoustic Context on Cortical Discrimination Responses

Experiment 1 investigated whether previously reported carrier- and direction-dependent ACC effects could be associated with context-dependent and rate-dependent influences on cortical discrimination responses. Previous work reported larger ACCs for tonal than noise carriers and stronger responses for transitions from an unmodulated reference to a spectrotemporally modulated target than for the reverse transition (Simonsen et al., 2025).

To examine these effects, ACCs and ASSRs were recorded from five NH participants using two classes of stimuli: continuous STMs and DSTMs. Stimuli were presented using either a TC or a NC. Responses were analysed separately for RT transitions, TR transitions, and the combination of both transition directions (R&T). For DSTM stimuli, the spectral transition corresponded to a 90° phase shift in the ripple pattern (DSTM-90).

To evaluate the possible contribution of response-rate adaptation, recordings were obtained using alternating periods of 2.0 and 3.3 s. Primary outcome measures were ACC response strength (F-value), N1-P2 amplitude, and N1 latency. These measures were compared across stimulus types, carrier conditions, and transition directions. ASSRs were analysed simultaneously to determine whether effects observed for the ACC could be accompanied by differences in phase-locked encoding of the ongoing stimulus envelope.

#### 2.7.2. Experiment 2: Dissociating Spectral and Intensity Contributions to Cortical Discrimination

Experiment 2 examined whether temporal-envelope variability could selectively alter intensity-driven and spectrally driven ACC responses.

Recordings were obtained from 12 NH listeners (two participants had also participated in Experiment 1). Three stimulus conditions were evaluated:

1. Intensity-only condition, in which transitions consisted exclusively of a 5-dB SPL intensity change (65-70 dB SPL or 70-65 dB SPL) without a change in spectral content.
2. Spectral-cue condition, in which transitions were generated using the DSTM-90 stimulus whilst maintaining overall stimulus level.
3. Combined-cue condition, in which spectral and intensity cues were presented simultaneously.

Each condition was presented with and without temporal roving. Responses were analysed separately for RT transitions, TR transitions, and the combination of both transition directions (R&T).

This experiment was designed to determine whether increasing temporal-envelope variability selectively reduces cortical responses elicited by intensity cues whilst preserving responses elicited by spectral changes. Outcome measures included ACC response strength, N1-P2 amplitude, response latency, and response detection rate.

#### 2.7.3. Experiment 3: Cortical Sensitivity to Spectral Contrast in Normal-Hearing and Hearing-Impaired Listeners

Experiment 3 characterised cortical representations of spectral contrast in NH and HI listeners and examined how these representations were influenced by temporal-envelope variability.

ACCs and ASSRs were recorded from 10 NH and 28 HI participants using the TC DSTM-90 stimulus. Of the NH listeners, four had participated in Experiment 2 and five in Experiment 1; two of these listeners had participated in both Experiments 1 and 2. Spectral contrast was manipulated by varying the modulation depth. Four CLs were tested: 0 (fully modulated), -4, -8, and −12 dB.

These values corresponded to ACT nCLs ranging from 16 dB nCL (highest contrast) to 4 dB nCL (lowest contrast).

Each contrast condition was measured both with and without temporal roving. Primary outcome measures included ACC response strength, N1-P2 amplitude, N1 latency, detection rate, and residual noise estimates. ASSRs were analysed in parallel to assess whether effects of spectral contrast on the ACC were accompanied by corresponding changes in phase-locked encoding of the 42-Hz stimulus envelope.

#### 2.7.4. Control Experiment: Effects of Absolute Presentation Level

A control experiment was performed to determine whether the differences observed between NH and HI listeners could be explained solely by differences in stimulus audibility.

ACCs and ASSRs were recorded using the TC DSTM-90 stimulus at 0 dB CL presented at either 65 or 50 dB SPL in 10 NH listeners. Measurements were obtained both with and without temporal roving.

This experiment was designed to determine whether the group differences observed between NH and HI listeners could be explained by reduced sensation level alone.

### 2.8. Statistical Analysis

All statistical analyses were performed in R software (R Development Core Team, 2026). Generalised linear mixed-effects (GLME) models were used to evaluate the effects of experimental factors on ACC and ASSR outcome measures. Separate models were fitted for response strength (F-value), amplitude, latency, and residual noise where appropriate. Participant was included as a random effect in all models whilst fixed effects varied across experiments.

Amplitude was modelled using a Gaussian error distribution, whereas F-values and latencies were modelled using Gamma distributions with logarithmic links.

Model reduction was performed through backward elimination of non-significant fixed-effect terms, with significance assessed using Wald *χ*^2^ tests (Wald, 1943; Fox and Weisberg, 2019). Following each reduction step, models were refitted and reassessed.

Model assumptions were evaluated through visual inspection of residuals and formal residual diagnostics using simulation-based approaches (Hartig, 2026, 5000 simulations). Outliers identified using a 1.5 interquartile-range criterion represented fewer than 2% of observations and were excluded prior to model refitting. Post hoc comparisons were performed using estimated marginal means with false discovery rate (FDR) correction for multiple comparisons.

To examine the association between cortical discrimination responses and behavioural performance, linear regression analyses were performed using ACT values as the dependent variable.

For each participant, ACC-derived predictors included: F-values, N1-P2 amplitudes, N1 latencies, slopes of the above measures across contrast levels, and differences between roving and non-roving conditions.

Initial models examined the relationship between individual ACC features and ACT values.

Additional models evaluated whether predictive performance improved when pure-tone average (PTA) of the better ear and participant age were included as predictors. Full models initially included the ACC feature, better-ear PTA, age, and their interactions. Backward stepwise reduction was then performed by sequentially comparing each model with a simpler nested model using an F-test. A term was retained when its removal significantly reduced explained variance. Thus, the contribution of the ACC feature was evaluated within the same model-reduction procedure while accounting for better-ear PTA, age, and retained interaction terms.

Model performance was quantified using the proportion of explained variance (*R*^2^). Statistical significance was assessed using a significance criterion of *α* = 0.05.

## Results

### 3.1. Cortical discrimination responses are modulated by spectrotemporal variance

In Experiment 1, we investigated whether differences in ACC responses previously reported by Simonsen et al. (2025) could be explained by context-dependent effects. ACCs were recorded in five NH participants using two carrier types, a NC and a TC, and two stimuli, STM and DSTM.

ACCs elicited by the TC (Figure 3a, bottom panels) exhibited larger response magnitudes than those elicited by the NC (Figure 3a, top panels), particularly for the DSTM-90 condition. Furthermore, ACCs evoked by the STM stimulus differed according to the direction of the acoustic transition. Responses to TR were reduced relative to RT changes (Figure 3, right panel), as reflected by lower response strength (F-value), smaller N1-P2 amplitudes, and longer N1 latencies (Figure 3b-d). This asymmetry was observed for both carrier types.

**Figure 3.**
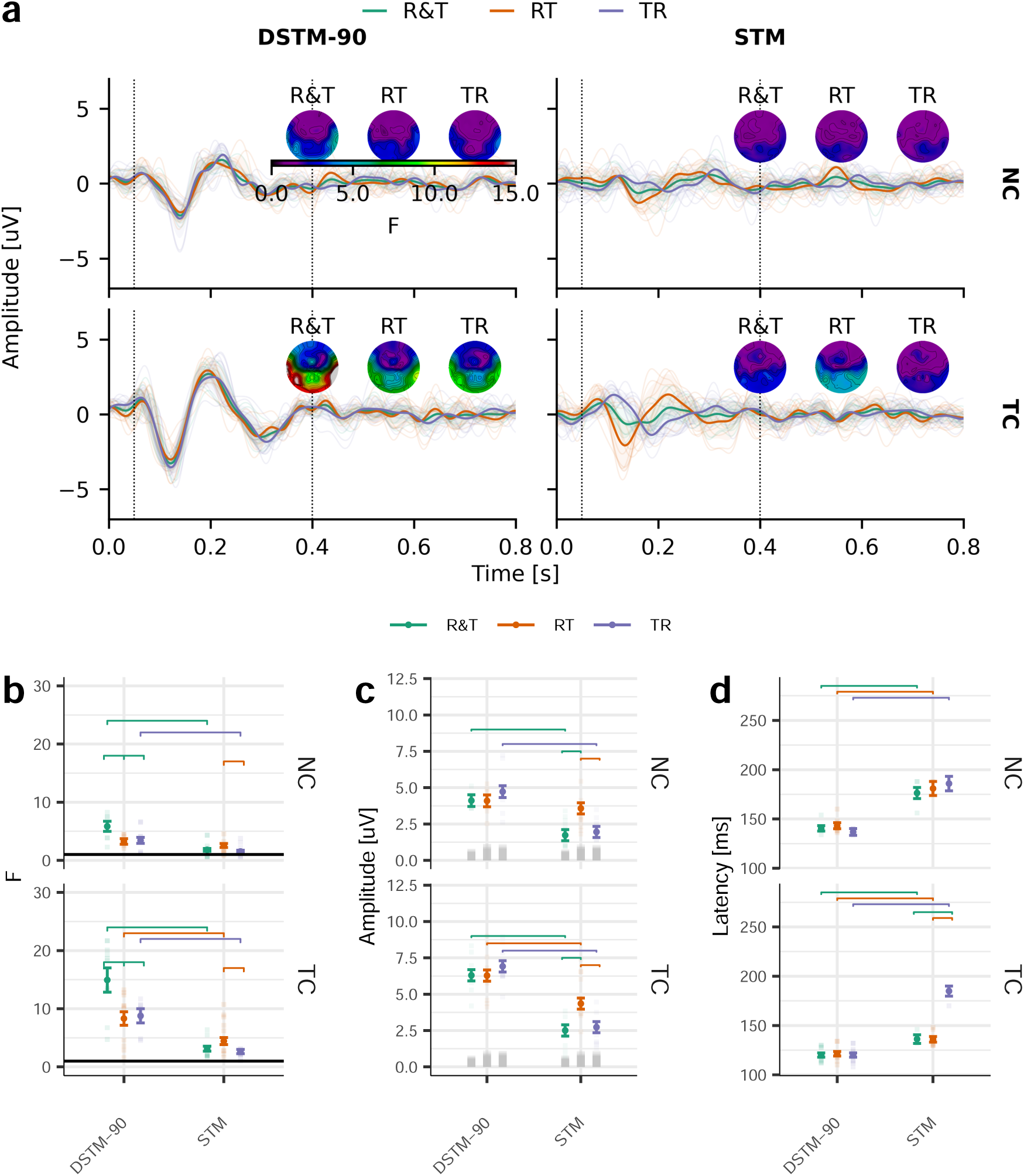
Effects of carrier type and transition direction on the ACC. **a**, ACC responses evoked by DSTM-90 (left) and STM (right) stimuli obtained from RT, TR, or both R&T transition conditions, indicated by the corresponding colour. The top panels show responses obtained using an NC, and the bottom panels show responses obtained using a TC. Individual responses are shown by the fine semitransparent traces (left and right mastoids) and averaged responses by the solid coloured traces. Topographic maps show the response strength (F-value) across the entire scalp for each transition direction. The vertical dashed lines indicate the analysis time window employed to compute the F-value. **b**, Response strength, **c** N1-P2 amplitudes, and **d** N1 latencies from the left and right mastoid electrodes. The stimulus is indicated on the horizontal axis and the carrier on the right side of the panels. Individual data are shown by the semitransparent points, whilst the mean and standard error of the GLME model are shown by the solid points and error bars respectively. Significant post hoc comparisons are shown by the horizontal bars within each panel (FDR corrected). Horizontal solid black lines in **b** indicate the unitary F-value (−∞ dB SNR). The semitransparent grey bars in **c** denote the residual noise (RN) in the responses.

To quantify these effects, GLME models were fitted with response strength (F-value), ACC amplitude, or N1 latency as dependent variables, and *stimulus type* (DSTM, STM), *carrier* (TC, NC), *transition direction* (RT, TR, or the combination of both transition directions R&T), and *recording channel* (left and right mastoid) as fixed effects. Significant main effects of *stimulus type* were observed for response strength [*χ*^2^(1) = 37.1, p < 0.001], amplitude [*χ*^2^(1) = 34.5, p < 0.001], and latency [*χ*^2^(1) = 38.1, p < 0.001].

Carrier type also significantly affected response strength [F-value: *χ*^2^(1) = 45.4, p < 0.001], amplitude [*χ*^2^(1) = 55.7, p < 0.001], and latency [*χ*^2^(1) = 37.4, p < 0.001]. A significant main effect of *transition direction* was observed for response strength [*χ*^2^(2) = 14.8, p < 0.001]. Furthermore, significant interactions were found between *stimulus type* and *transition direction* for response strength [*χ*^2^(2) = 16.6, p < 0.001] and amplitude [*χ*^2^(2) = 24.7, p < 0.001]. Significant interactions between *stimulus type* and *carrier* were also observed for response strength [*χ*^2^(2) = 4.1, p = 0.042] and amplitude [*χ*^2^(2) = 12.5, p < 0.001], and a significant three-way interaction between *stimulus type*, *carrier*, and *transition direction* was observed for the N1 latency [*χ*^2^(4) = 66.7, p < 0.001].

These interactions are illustrated in Figure 3b-d. For the DSTM stimulus, RT and TR transitions produced comparable ACC responses. Consequently, averaging responses across both transition directions (R&T) increased the SNR, resulting in larger F-values. In contrast, for the STM stimulus, RT transitions evoked significantly larger responses than TR transitions. Averaging across directions therefore reduced overall response strength, yielding lower F-values in the combined R&T condition.

Similar analyses were performed for ASSRs (Sup. Figure 1a-d). Significant main effects of *carrier* were observed for response strength (F-value) [*χ*^2^(1) = 42.4, p < 0.001; Sup. Figure 1b], amplitude [*χ*^2^(1) = 44.8, p < 0.001; Sup. Figure 1c] and latency [*χ*^2^(1) = 17.7, p < 0.001; Sup. Figure 1d]. Response strength was significantly affected by *transition direction* [*χ*^2^(2) = 41.7, p < 0.001; Sup. Figure 1b], whilst recording *channel* significantly influenced both amplitude [*χ*^2^(1) = 10.0, p = 0.002] and latency [*χ*^2^(1) = 10.1, p = 0.002]. A significant interaction between *stimulus type* and *transition direction* was observed for latency [*χ*^2^(2) = 23.2, p < 0.001], and a significant interaction between *stimulus type* and *carrier* was found for response strength [*χ*^2^(1) = 20.4, p < 0.001] and amplitude [*χ*^2^(1) = 12.3, p < 0.001]. Overall,

ASSR amplitudes were similar for RT and TR transition blocks, such that combining responses across transition directions improved SNR. The *stimulus type* × *carrier* interaction was driven by larger amplitudes for the DSTM than STM stimulus when presented with the TC. In contrast, no significant amplitude differences between DSTM and STM were observed for the NC. Likewise, ASSR amplitudes evoked by STM stimuli were comparable across carrier types.

### 3.2. Cortical differences between stimuli are not explained by rate adaptation

To determine whether the smaller ACCs observed for the STM stimulus and the NC condition could be explained by rate adaptation—specifically, the presentation rate of the transitions evoking the ACC—ACCs were measured using a longer alternating period of 3.3 s in Experiment 1 (Figure 4).

**Figure 4.**
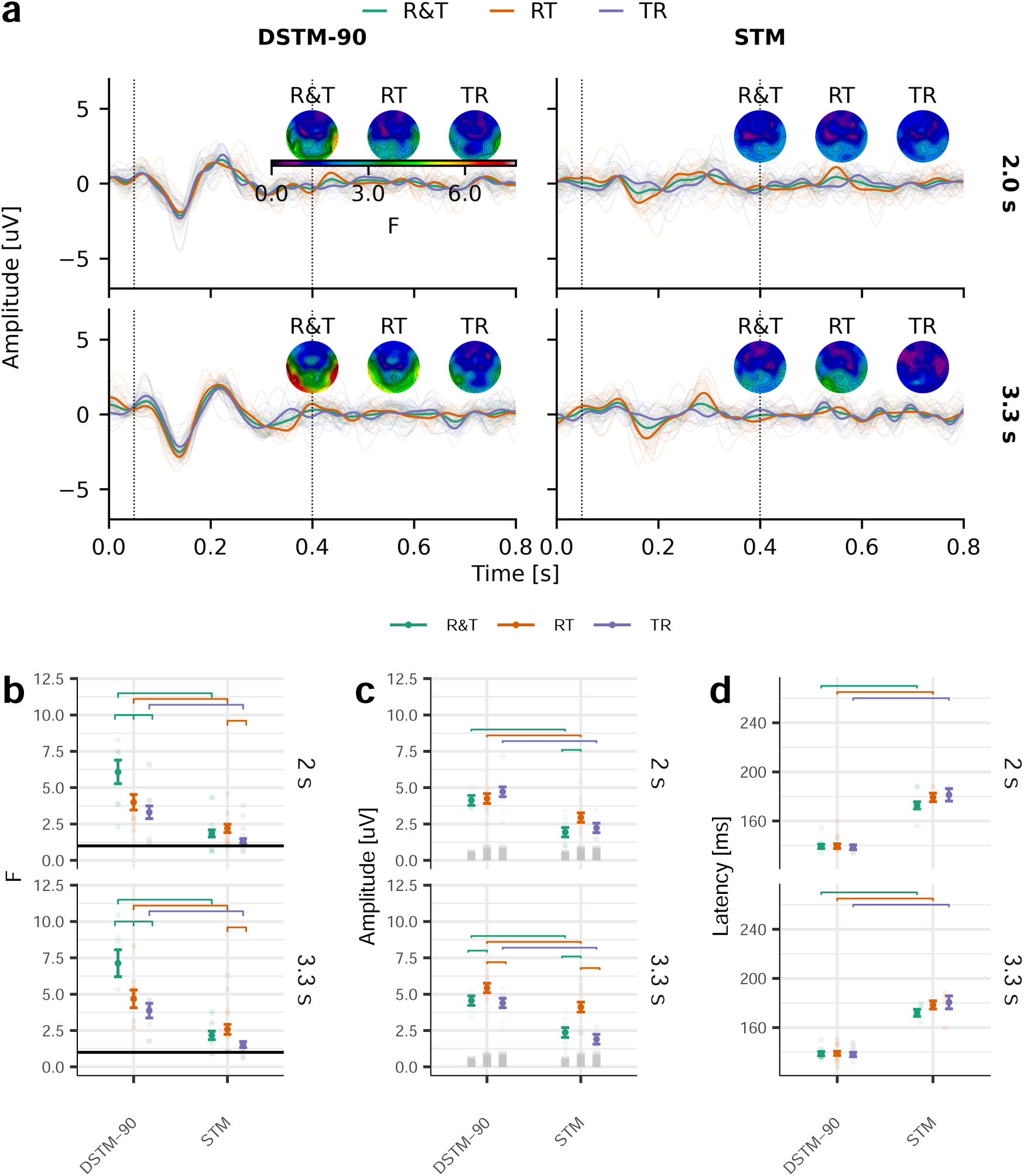
Effects of alternating period on ACC responses. **a**, ACC responses to DSTM-90 (left) and STM (right) obtained by changes from RT, TR, or both R&T, indicated by the corresponding colour. Responses were recorded using the NC. The top panels show responses obtained using a 2.0-s alternation period, whereas the bottom panels show responses obtained using a 3.3-s alternation period. Individual responses are shown by the fine semitransparent traces (left and right mastoids) and averaged responses by the solid coloured traces. Topographic maps show the response strength (F-value) across the entire scalp for each transition direction. The vertical dashed lines indicate the analysis time window employed to compute the F-value. **b**, Response strength, **c** N1-P2 amplitudes, and **d** N1 latencies from the left and right mastoid electrodes. The stimulus is indicated on the horizontal axis and the alternating period on the right side of the panels. Individual data are shown by the semitransparent points, whilst the mean and standard error of the GLME model are shown by the solid points and error bars respectively. Significant post hoc comparisons are shown by the horizontal bars within each panel (FDR corrected). Horizontal solid black lines in **b** indicate the unitary F-value (−∞ dB SNR). The semitransparent grey bars in **c** denote the RN in the responses.

Consistent with reduced rate adaptation, increasing the alternating period from 2.0 s to 3.3 s generally increased ACC response strength. However, the overall pattern of results remained similar across alternating periods (cf. Figure 4 at 2.0 and 3.3 s). Specifically, the differences between STM and DSTM stimuli, as well as the directional asymmetries observed for STM responses, were preserved at the longer interval.

To quantify these effects, GLME models were fitted with response strength (F-value), amplitude, or latency as dependent variables, and stimulus type (STM, DSTM), alternating period (2.0 and 3.3 s), transition direction (RT, TR, and R&T), and recording channel (left and right mastoid) as fixed effects.

Significant interactions were observed between *alternating period* and *transition direction* for both response strength [*χ*^2^(2) = 9.9, p = 0.007] and amplitude [*χ*^2^(2) = 14.1, p < 0.001]. Significant interactions between *stimulus type* and *transition direction* were also identified for response strength [*χ*^2^(2) = 9.2, p = 0.010] and amplitude [*χ*^2^(2) = 9.3, p = 0.009]. In addition, significant main effects of *transition direction* were found for response strength [*χ*^2^(2) = 8.7, p = 0.013], whilst stimulus type significantly affected response strength [*χ*^2^(1) = 63.9, p < 0.001], amplitude [*χ*^2^(1) = 60.1, p < 0.001], and latency [*χ*^2^(1) = 347.5, p < 0.001].

Comparison of the two alternating periods revealed significantly larger F-values at 3.3 s than at 2.0 s for RT transitions, corresponding to an increase of approximately 35% for both DSTM-90 and STM stimuli. Nevertheless, the increase associated with the longer interval did not account for the differences previously observed between carrier types. For example, the response strength obtained for the R&T condition using the DSTM-90 stimulus with a TC at a 2.0 s alternating period remained approximately twice that obtained for the same stimulus presented with a NC at a 3.3 s alternating period [F-values: 14.94 vs. 7.35; cf. Figure 3b and Figure 4b]. Similarly, for STM stimuli, RT responses elicited with a TC at a 2.0 s alternating period were approximately 50% larger than those elicited with a NC at a 3.3 s alternating period [F-values: 4.44 vs. 2.99]. These differences were also reflected in response amplitudes [Figure 4c].

Response latencies differed markedly between stimulus types, with STM stimuli producing significantly longer latencies than DSTM stimuli. In contrast, latency measures were not significantly affected by alternating period (Figure 4d). Furthermore, the effects of alternating period on response strength could not be attributed to changes in residual noise levels, as no significant effect of alternating period was observed in the corresponding GLME analysis (all p > 0.05).

Taken together, these findings indicate that although extending the alternating period enhances ACC response strength, the effects of carrier type and stimulus structure persist. In particular, the reduced responses observed for NC relative to TC conditions and the directional asymmetry associated with STM stimuli were maintained at the longer interval, suggesting that these effects cannot be explained solely by rate adaptation.

### 3.3. Temporal-envelope variability partially dissociates intensity and spectral contributions to cortical responses

In Experiment 2, we examined whether the potential contribution of intensity cues to ACC generation could be reduced by increasing temporal-envelope variability. To this end, ACCs were recorded in 12 participants using a TC DSTM-90-like stimulus containing a 5 dB SPL change but no spectral change. In this paradigm, RT transitions corresponded to intensity increments (65 to 70 dB SPL), whereas TR transitions corresponded to intensity decrements (70 to 65 dB SPL). Responses were measured both in the presence and absence of a second-order speech-like temporal modulation (roving) and were compared with ACCs elicited by DSTM-90 stimuli containing a spectral cue, presented either with or without concurrent intensity cues (Figure 5).

**Figure 5.**
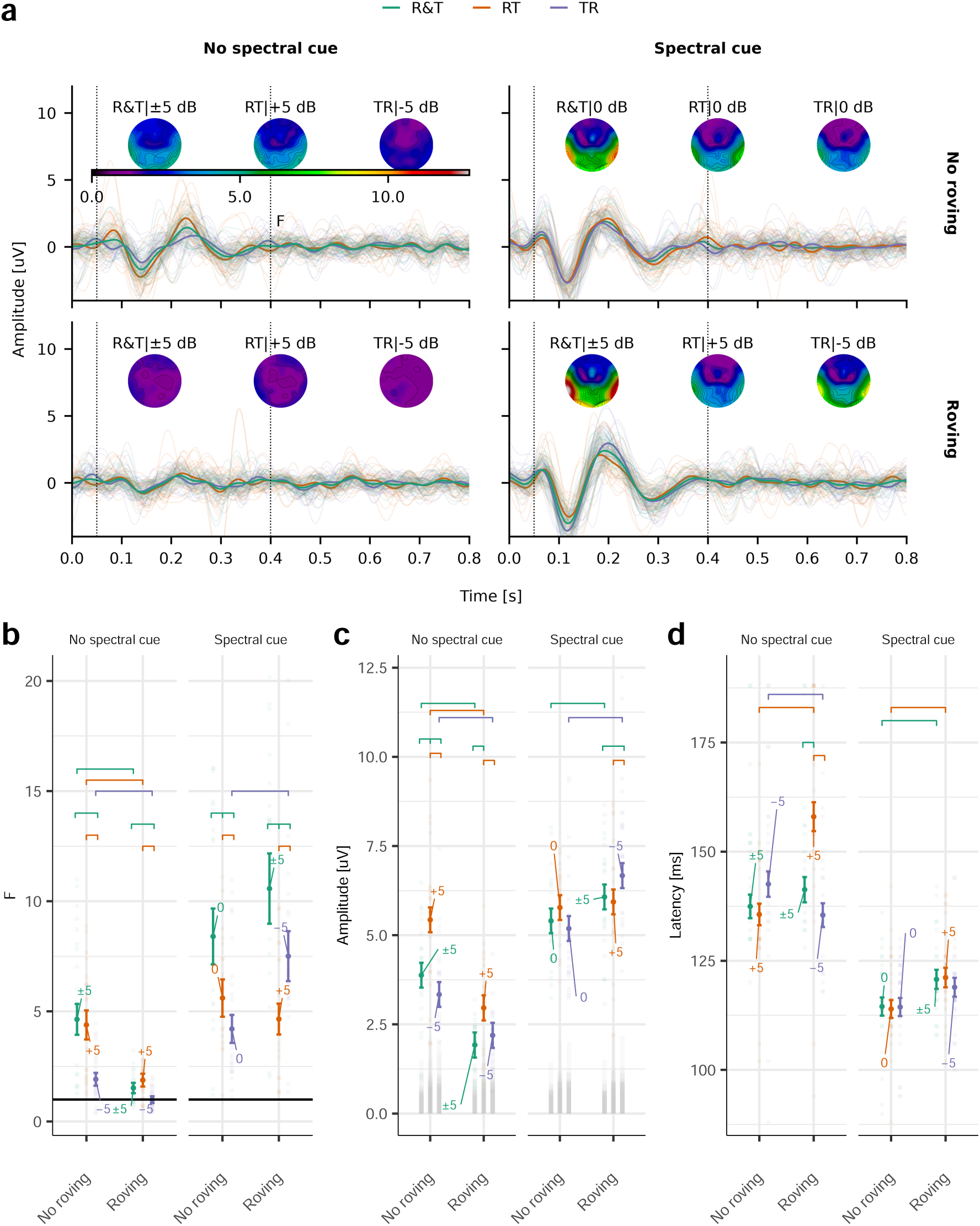
Dissociating loudness-related and spectral contributions to the ACC. **a**, ACC responses to TC DSTM-90-like stimuli, in the absence of spectral changes (no spectral cue, left) and to a TC DSTM-90 (spectral cue, right). Responses were obtained for RT, TR, or both R&T transition directions, indicated by the corresponding colour, at an alternating period of 2.0 s. The top panels show ACCs obtained without temporal roving, whereas the bottom panels show ACCs obtained with temporal roving. Individual responses are shown by the fine semitransparent traces (left and right mastoids) and averaged responses by the solid coloured traces. Topographic maps show the response strength (F-value) across the entire scalp for each transition direction. The difference in stimulus intensity included in the transition-direction averages is indicated on top of the topographic maps: 65 to 70 dB SPL: +5 dB; 70 to 65 dB SPL: −5 dB; 65 to 65 dB SPL: 0 dB; and both 65 to 70 dB SPL and 70 to 65 dB SPL: ±5 dB. The vertical dashed lines indicate the analysis time window employed to compute the F-value. **b**, Response strength, **c** N1-P2 amplitudes, and **d** N1 latencies from the left and right mastoid electrodes. The temporal roving condition is indicated on the horizontal axis. Individual data are shown by the semitransparent points, whilst the mean and standard error of the GLME model are shown by the solid points and error bars respectively. Significant post hoc comparisons are shown by the horizontal bars within each panel (FDR corrected). Horizontal solid black lines in **b** indicate the unitary F-value (−∞ dB SNR). The semitransparent grey bars in **c** denote the RN in the responses. The coloured numbers indicate the intensity difference, in dB, for each transition direction.

In the absence of spectral changes, intensity transitions elicited robust ACCs in both directions, although responses to intensity decrements were generally smaller than responses to increments (Figure 5a). Introducing temporal roving substantially reduced response strength in the intensity-only condition. In contrast, ACCs elicited by stimuli containing spectral cues did not decrease with roving, exhibiting comparable waveform morphology and numerically larger responses in the roving than in the no-roving condition (Figure 5a).

Response detection rates showed a similar pattern. For the R&T condition, which produced the highest response strength in the intensity-only paradigm, detection rates averaged 75.0% across mastoid electrodes in the absence of roving but decreased to 18.2% when roving was introduced. In contrast, when a spectral cue was present, detection rates remained high both without roving (91.7%) and with roving (95.8%).

These observations were supported by GLME analyses of response strength (F-value), N1-P2 amplitude, and latency. Models included *spectral cue* (present or absent), *roving* condition (present or absent), *transition direction* (RT, TR, and R&T), and *recording channel* as fixed effects. Significant three-way interactions were observed between *roving*, *spectral cue*, and *transition direction* for response strength [*χ*^2^(2) = 8.4, p = 0.015] and latency [*χ*^2^(2) = 18.2, p < 0.001].

Significant two-way interactions were also identified between *roving* and *transition direction* [amplitude: *χ*^2^(2) = 24.2, p < 0.001; latency: *χ*^2^(2) = 40.2, p < 0.001], *roving* and *spectral cue* [F-value: *χ*^2^(1) = 51.1, p < 0.001; amplitude: *χ*^2^(1) = 141.2, p < 0.001], and *transition direction* and *spectral cue* [F-value: *χ*^2^(2) = 8.7, p = 0.013; amplitude: *χ*^2^(2) = 34.3, p < 0.001]. Significant main effects were observed for *roving*, *spectral cue*, and *transition direction* across multiple response measures.

The interaction between roving and spectral cue was particularly evident in the response-strength measures (Figure 5b). In the intensity-only condition, averaging across transition directions improved the signal-to-noise ratio when roving was absent, resulting in the largest F-values for the R&T condition. Temporal roving substantially reduced response strength and altered this pattern, with responses becoming dominated by intensity increments. As a consequence, the R&T response was markedly diminished, with F-values approximately 2.3-fold lower than those observed without roving.

In contrast, when a spectral cue was present, the R&T condition consistently produced the strongest responses regardless of whether temporal roving was applied. Both the individual participant data and model-derived means showed numerically greater response strength under the roving condition, as reflected in the F-values and amplitudes (Figure 5b&c), indicating that the effects of temporal-envelope variability differed markedly between intensity-driven and spectrally driven cortical discrimination responses. Response amplitudes exhibited a partially different pattern (Figure 5c). In the absence of spectral cues, amplitude estimates were broadly comparable across roving conditions despite the substantial reduction in response strength. Moreover, RT amplitudes measured in the intensity-only condition were similar to those observed when spectral cues were available in the absence of roving. This finding suggests that even when nominal SPLs are matched, perceptual loudness differences arising from variations in spectral content and spectral weighting could contribute to the observed ACC amplitudes.

Response latencies revealed the clearest separation between intensity- and spectral-driven responses (Figure 5d). Conditions containing a spectral cue produced shorter and more consistent latencies across transition directions than conditions relying exclusively on intensity cues. By contrast, intensity-only responses exhibited longer latencies and greater variability, particularly in the presence of temporal roving. These latency differences were reflected in the significant interactions involving spectral cue, roving condition, and transition direction observed in the statistical analyses.

Collectively, these findings demonstrate that increasing temporal-envelope variability strongly attenuates ACCs elicited by intensity changes whilst preserving, and in some conditions slightly enhancing, responses elicited by spectral changes in NH listeners. The persistence of robust ACCs in the spectral-cue conditions, despite the marked reduction of intensity-driven responses, indicates that the contributions of intensity and spectral cues to cortical discrimination can be at least partially dissociated. Furthermore, the shorter and more consistent response latencies observed in the spectral-cue conditions suggest that cortical discrimination was more strongly driven by spectral information when such cues were available.

### 3.4. Cortical sensitivity to spectral contrast in NH and HI participants

Having established that ACCs elicited by the TC DSTM stimulus were robust and that speech-like temporal roving effectively reduced contributions from potential loudness cues, Experiment 3 examined cortical sensitivity to spectral contrast in NH and HI participants. ACCs were elicited using a TC DSTM-90 stimulus across a range of modulation levels, here referred to as CLs, from 0 dB CL—corresponding to a fully modulated and perceptually most salient stimulus—to −12 dB CL—corresponding to the least salient stimulus tested here—in 4-dB steps. A total of 10 NH and 28 HI participants were assessed under both roving and no-roving conditions (Figure 6). ACC responses decreased as spectral contrast was reduced and were consistently smaller in HI than in NH listeners (Figure 6a&b). In NH participants, response strength did not decrease monotonically with CL in the absence of temporal roving. Instead, responses at the highest contrast level were comparable to, or slightly smaller than, those observed at intermediate contrast levels (Figure 6c). This non-monotonic pattern disappeared when temporal roving was introduced, revealing a systematic reduction in ACC strength with decreasing CL. Consistent with the findings of Experiment 2, NH listeners exhibited numerically larger responses at 0 dB CL in the roving condition than in the no-roving condition.

**Figure 6.**
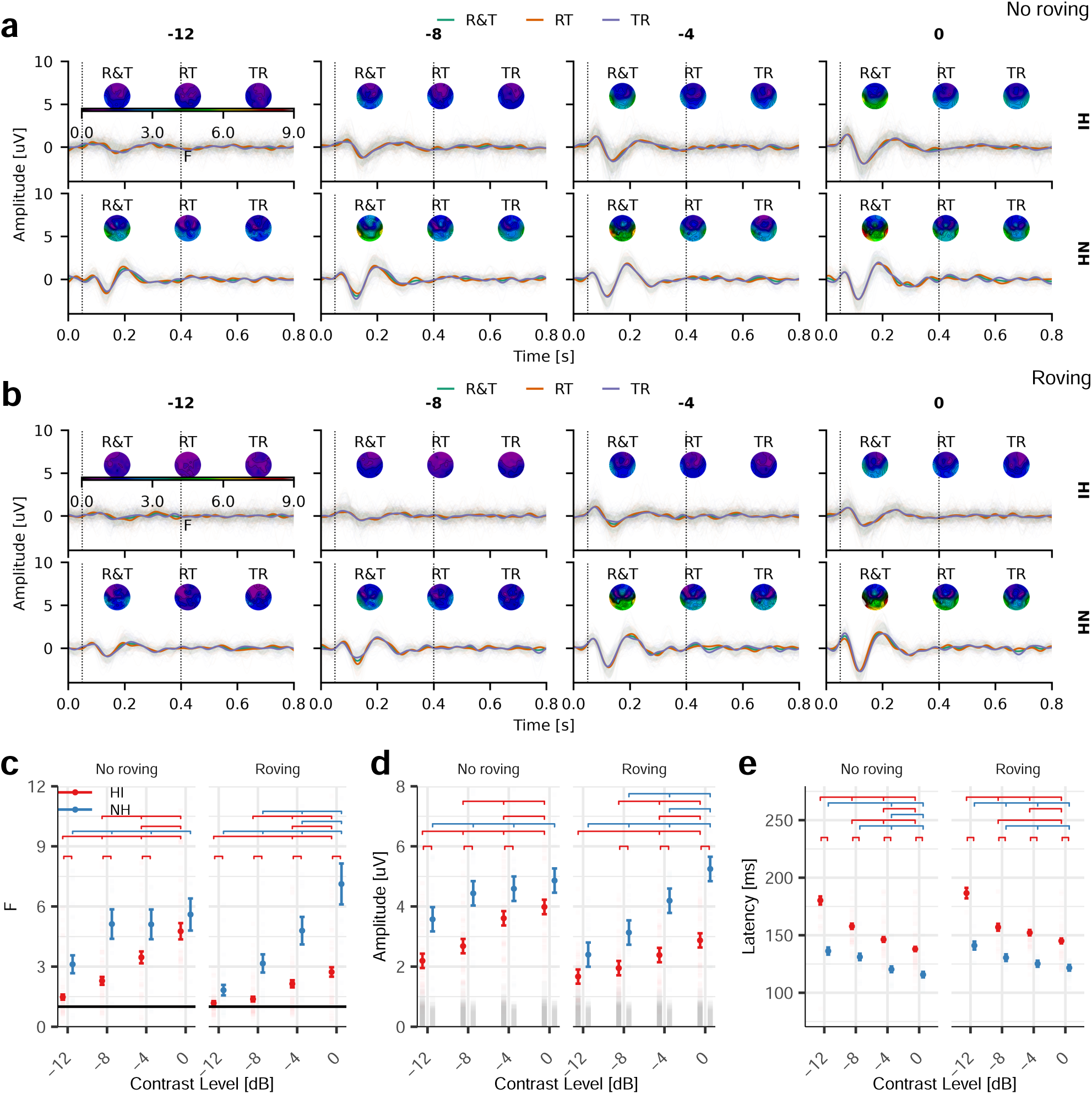
Effects of spectral contrast and temporal roving on ACCs in NH and HI. **a**, ACC responses evoked by DSTM-90 stimuli using a TC without temporal roving. RT, TR, and combined R&T transitions are indicated by the corresponding colour. Responses are shown for HI listeners (top) and NH listeners (bottom). The spectral CL is indicated above each panel. Individual responses are shown as semitransparent traces (left and right mastoids) and grand-average responses by solid traces. Topographic maps show the average response strength (F-value) across the scalp. Vertical dashed lines indicate the analysis window. **b**, Same as panel **a**, but with temporal roving applied. **c**, Response strength, **d** N1-P2 amplitude, and **e** N1 latency. Data in **c**-**e** represent R&T responses averaged across left and right mastoid electrodes. Individual observations are shown as semitransparent points, whilst solid symbols and error bars represent estimated marginal means ± standard error from the GLME. Significant post hoc comparisons (FDR corrected) are indicated by horizontal bars. Temporal roving produced a more monotonic contrast-response function in NH listeners whilst reducing response strength and increasing group separation between NH and HI listeners. The horizontal solid black line in **c** indicates the unitary F-value (−∞ dB SNR). Semitransparent grey bars in **d** denote RN.

In contrast, HI listeners showed a monotonic reduction in response strength with decreasing CL under both roving conditions. Temporal roving produced a pronounced reduction in ACC magnitude across all contrast levels. At 0 dB CL, response strength in HI listeners was reduced by more than two-thirds relative to the no-roving condition and was less than half that observed in NH listeners under equivalent stimulus conditions.

These observations were confirmed by GLME analyses of response strength (F-value), N1-P2 amplitude, and N1 latency. Models included CL, *roving* condition, *hearing group*, and *recording channel* as fixed effects. Significant main effects of CL were observed for both F-values [*χ*^2^(3) = 237.4, p < 0.001] and amplitudes [*χ*^2^(3) = 180.5, p < 0.001], indicating progressively weaker cortical responses with decreasing spectral contrast. Significant main effects of roving (F-value: *χ*^2^(1) = 8.0, p = 0.005; amplitude: *χ*^2^(1) = 12.6, p < 0.001) and hearing group (F-value: *χ*^2^(1) = 20.1, p < 0.001; amplitude: *χ*^2^(1) = 8.7, p = 0.003) were also observed.

Importantly, these effects depended on both hearing status and stimulus contrast. Significant interactions were found between CL and *roving* [F-value: *χ*^2^(3) = 10.2, p = 0.017; amplitude: *χ*^2^(3) = 14.5, p = 0.002], CL and *hearing group* [F-value: *χ*^2^(3) = 22.4, p < 0.001; amplitude: *χ*^2^(3) = 11.0, p = 0.012], and *roving* and *hearing group* [F-value: *χ*^2^(1) = 3.9, p = 0.048; amplitude: *χ*^2^(1) = 5.1, p = 0.025]. Furthermore, significant three-way interactions between CL, *roving*, and *hearing group* were observed for both F-values [*χ*^2^(3) = 28.0, p < 0.001] and amplitudes [*χ*^2^(3) = 39.1, p < 0.001], indicating that the influence of temporal-envelope variability differed between NH and HI listeners as a function of spectral contrast.

Analysis of response latencies revealed significant effects of CL [*χ*^2^(3) = 240.7, p < 0.001] and *hearing group* [*χ*^2^(1) = 96.1, p < 0.001], as well as significant interactions between CL and *hearing group* [*χ*^2^(3) = 20.4, p < 0.001] and between *CL* and *roving* [*χ*^2^(3) = 7.9, p = 0.048]. Across both listener groups, response latencies increased systematically as spectral contrast decreased.

Post hoc analyses indicated that, among NH listeners, temporal roving was associated with slightly larger F-values at 0 dB CL, although this effect did not reach statistical significance [z = 1.53, p = 0.080]. At lower contrast levels (-8 and −12 dB CL), however, both response strength and amplitude were significantly reduced by temporal roving (all p < 0.01). In HI listeners, temporal roving significantly reduced F-values and amplitudes at all contrast levels examined [all p < 0.001]. Direct comparisons between hearing groups revealed that, in the absence of roving, NH and HI listeners produced comparable response strengths at 0 dB CL. Group differences emerged as contrast decreased, reflecting a steeper decline in cortical response magnitude in HI listeners. When temporal roving was introduced, NH and HI groups differed significantly across all contrast levels for F-values (all p ≤ 0.021), and across all but the lowest contrast condition for amplitudes (all p ≤ 0.015 for CLs ≥ −8 dB).

Response latency provided an additional source of group separation. Latencies were consistently longer in HI than NH listeners and increased as spectral contrast decreased. Mean N1 latencies ranged from approximately 116 to 141 ms in NH participants and from 138 to 186 ms in HI participants. Temporal roving further increased response latencies at 0 and −4 dB CLs in both groups, producing an average prolongation of approximately 6 ms [all p < 0.01].

These effects were also reflected in ACC detection rates. In NH listeners, detection rates at 0 dB CLs increased from 85% in the no-roving condition to 95% when temporal roving was introduced. In contrast, HI listeners showed a marked decline in detection rates under roving conditions, decreasing from 93% to 59%.

ASSR results (Sup. Figure 2) showed a different pattern. ASSR F-values (R&T) were significantly larger in NH than in HI listeners [*χ*^2^(1) = 23.6, p < 0.001] and were reduced by temporal roving [*χ*^2^(1) = 25.2, p < 0.001]. ASSR latencies were also shorter in NH than in HI listeners [*χ*^2^(1) = 6.9, p = 0.009], irrespective of temporal-roving condition. Residual noise levels were also significantly higher in HI participants (30.6 nV) than in NH participants (23.6 nV; *χ*^2^(1) = 8.6, p = 0.003). However, unlike the ACC measures, ASSR F-values, amplitudes, and latencies showed no systematic dependence on spectral contrast level. Consistent with this observation, ASSR detection rates remained near ceiling across all conditions (100% in NH listeners and 98.5% in HI listeners).

Together, these findings demonstrate that cortical sensitivity to spectral contrast declines with decreasing contrast and is substantially reduced in HI listeners. Temporal roving accentuates these group differences, particularly at high contrast levels, resulting in clearer separation between NH and HI participants. In contrast, ASSRs remained robust and showed no systematic dependence on spectral contrast. This dissociation suggests that the contrast-dependent ACC effects primarily reflect change-related cortical discrimination rather than a general loss of phase-locked envelope encoding.

### 3.5. Cortical discrimination responses remain robust across presentation levels

Because ASSR detection rates remained near ceiling in both hearing groups, an additional control experiment was conducted in NH participants to evaluate the influence of presentation level on ACC responses. ACCs and ASSRs were therefore recorded using the TC DSTM-90 stimulus (0 dB CL) presented at either 65 or 50 dB SPL, and responses were compared across presentation levels.

Despite the 15-dB reduction in stimulus level, ACCs remained readily detectable at 50 dB SPL (Figure 7). GLME analyses revealed significant effects of presentation level on both response strength [*χ*^2^(1) = 5.8, p = 0.016] and N1-P2 amplitude [*χ*^2^(1) = 7.7, p = 0.005]. In contrast, response latency was unaffected by presentation level, remaining stable at approximately 117 ms across conditions. Relative to the 65 dB SPL condition, average F-values and amplitudes at 50 dB SPL were reduced by approximately 18% and 11%, respectively.

**Figure 7.**
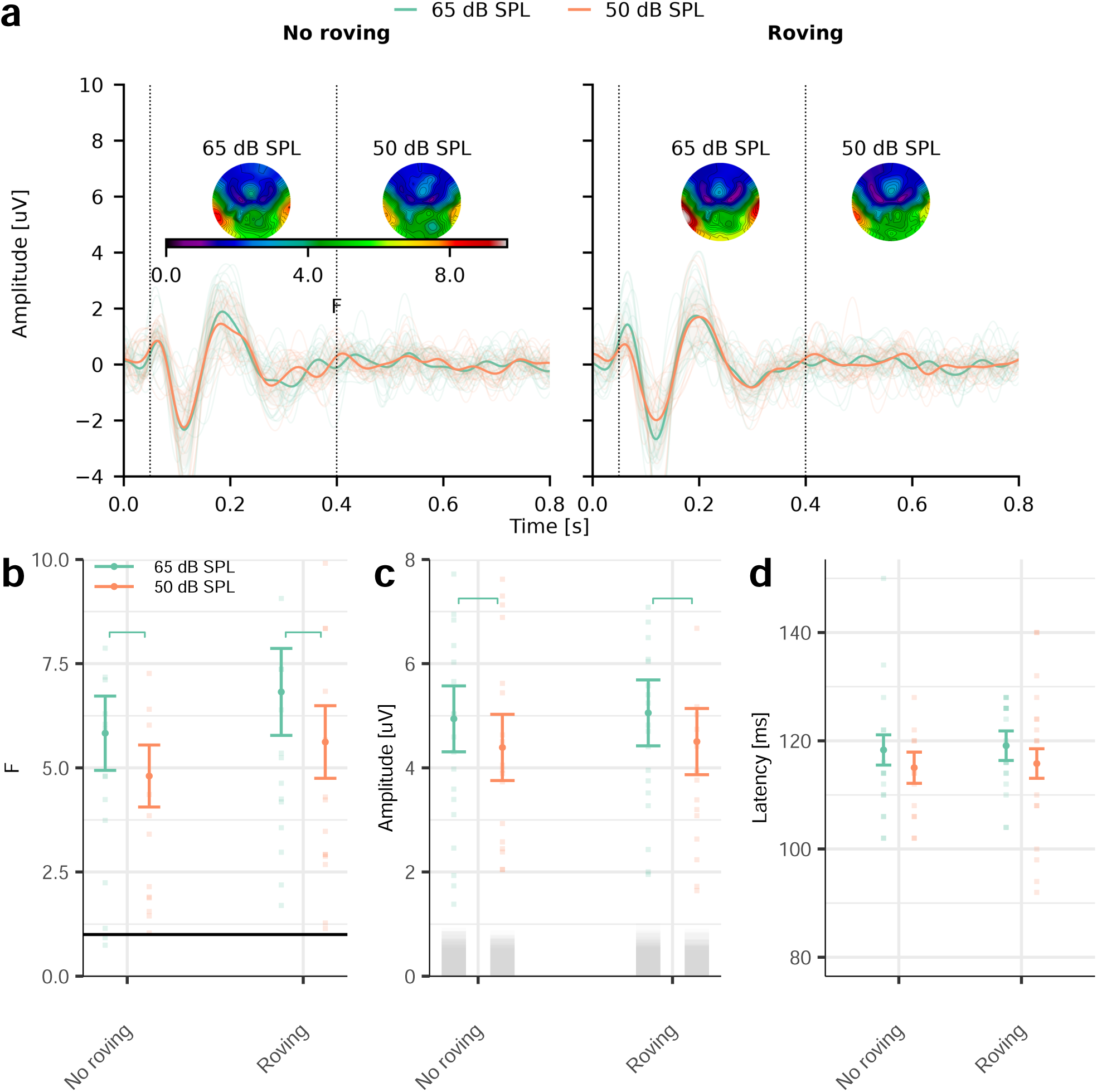
Effects of presentation level and temporal roving on ACC responses in NH listeners. **a**, ACC waveforms evoked by DSTM-90 using a TC at presentation levels of 50 dB SPL (orange) and 65 dB SPL (green), without temporal roving (left) and with temporal roving (right). Spectral contrast was fixed at 0 dB CL. Individual responses are shown as semitransparent traces (left and right mastoids) and grand-average responses as solid traces. Topographic maps illustrate mean ACC response strength (F-value) across the scalp. Vertical dashed lines indicate the analysis window. **b**, Response strength, **c** N1-P2 amplitude, and **d** N1 latency. Data in **b**-**d** represent R&T responses averaged across mastoid electrodes. Individual observations are shown as semitransparent points, whilst solid symbols and error bars represent estimated marginal means ± standard error from the GLME. Significant post hoc comparisons (FDR corrected) are indicated by horizontal bars. The horizontal black line in **b** indicates the unitary F-value (−∞ dB SNR). Semitransparent grey bars in **c** denote RN.

A similar pattern was observed for ACC detection rates. At 65 dB SPL, detection rates were 95% and 85% in the roving and no-roving conditions, respectively. Reducing the presentation level to 50 dB SPL resulted in only a modest decrease in detection rates, to 89% with roving and 72% without roving. Thus, although lower stimulus levels reduced ACC strength, responses remained robust and detectable in the majority of participants.

To place these effects in context, NH responses were compared with those obtained from HI listeners. Under roving conditions at 65 dB SPL, HI participants exhibited substantially smaller ACCs than NH participants, with response strength and amplitudes reduced by approximately 62% and 45%, respectively (Figure 6c&d). Importantly, these group differences persisted even when NH responses recorded at 50 dB were used as the reference. Compared with NH listeners tested at the lower presentation level, HI participants still showed reductions of approximately 48% in F-values and 36% in amplitudes under equivalent roving conditions.

ASSRs (Sup. Figure 3) exhibited a different pattern. Presentation level significantly affected ASSR response strength, amplitude, and latency, with lower SPLs producing weaker responses and longer latencies. Temporal roving also reduced ASSR F-values and amplitudes. Unlike the ACC, however, these changes were largely consistent with the effective reduction in stimulus level associated with the roving manipulation.

Taken together, these findings demonstrate that ACC responses exhibit only a modest dependence on absolute presentation level over the range tested. Although reducing stimulus level decreased response strength, the magnitude of this effect was substantially smaller than the differences observed between NH and HI listeners. The marked reduction in ACC responses observed in the HI group relative to the NH group in Experiment 3 therefore cannot be explained solely by reduced audibility and is more consistent with impaired cortical discrimination of the spectrotemporal contrast cues contained in the stimulus.

### 3.6. Cortical responses underlying discrimination are associated with behavioural performance

We next examined the relationship between ACC-derived measures and behavioural ACT values. To this end, a range of ACC features were extracted, including response strength (F-value), N1-P2 amplitude, and latency at individual CLs. In addition, linear slopes describing the relationship between log2-transformed ACC features and CL were calculated to quantify the rate at which cortical responses changed as spectral contrast decreased. These features were extracted separately for the roving and no-roving conditions, and differences between the two conditions were also considered to capture within participant roving effects.

The associations between individual ACC features and ACT values are summarised in Table 1 and Table 2.

**Table 1.** Associations between individual ACC features and ACT values for NH and HI participants. From left to right: Roving condition, ACC feature, explained variance (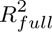), number of data points (N), and p values.

| Condition | ACC feature | $R_{full}^2$ | N | p |
| --- | --- | --- | --- | --- |
| No roving | 0 dB CL F | 0.07 | 46 | 0.08 |
| No roving | 0 dB CL Amplitude | 0.12 | 46 | 0.02 |
| No roving | 0 dB CL Latency | 0.41 | 41 | <0.01 |
| No roving | -4 dB CL F | 0.16 | 38 | 0.01 |
| No roving | -4 dB CL Amplitude | 0.10 | 38 | 0.05 |
| No roving | -4 dB CL Latency | 0.39 | 29 | <0.01 |
| No roving | -8 dB CL F | 0.25 | 38 | <0.01 |
| No roving | -8 dB CL Amplitude | 0.26 | 38 | <0.01 |
| No roving | -8 dB CL Latency | 0.54 | 25 | <0.01 |
| No roving | -12 dB CL F | 0.16 | 38 | 0.01 |
| No roving | -12 dB CL Amplitude | 0.14 | 38 | 0.02 |
| No roving | -12 dB CL Latency | 0.07 | 13 | 0.37 |
| Roving | 0 dB CL F | 0.38 | 46 | <0.01 |
| Roving | 0 dB CL Amplitude | 0.41 | 46 | <0.01 |
| Roving | 0 dB CL Latency | 0.48 | 34 | <0.01 |
| Roving | -4 dB CL F | 0.22 | 38 | <0.01 |
| Roving | -4 dB CL Amplitude | 0.33 | 38 | <0.01 |
| Roving | -4 dB CL Latency | 0.68 | 21 | <0.01 |
| Roving | -8 dB CL F | 0.28 | 38 | <0.01 |
| Roving | -8 dB CL Amplitude | 0.10 | 38 | 0.05 |
| Roving | -8 dB CL Latency | 0.41 | 15 | 0.01 |
| Roving | -12 dB CL F | 0.07 | 38 | 0.12 |
| Roving | -12 dB CL Amplitude | 0.03 | 38 | 0.31 |
| Roving | -12 dB CL Latency | 0.80 | 9 | <0.01 |
| Δ Roving | 0 dB CL F | 0.36 | 46 | <0.01 |
| Δ Roving | 0 dB CL Amplitude | 0.27 | 46 | <0.01 |
| Δ Roving | -4 dB CL F | 0.01 | 38 | 0.64 |
| Δ Roving | -4 dB CL Amplitude | 0.10 | 38 | 0.05 |
| Δ Roving | -8 dB CL F | 0.01 | 38 | 0.65 |
| Δ Roving | -8 dB CL Amplitude | 0.06 | 38 | 0.15 |
| Δ Roving | -12 dB CL F | 0.07 | 38 | 0.1 |
| Δ Roving | -12 dB CL Amplitude | 0.06 | 38 | 0.12 |
| No roving | F-value slope | 0.08 | 38 | 0.1 |
| No roving | Amplitude slope [log2/CL dB] | 0.03 | 38 | 0.29 |
| Roving | F-value slope | 0.25 | 38 | <0.01 |
| Roving | Amplitude slope [log2/CL dB] | 0.23 | 38 | <0.01 |
| Δ Roving | F slope [log2/CL dB] | 0.11 | 28 | 0.09 |
| Δ Roving | Amplitude slope [log2/CL dB] | 0.10 | 28 | 0.09 |

**Table 2.** Associations between individual ACC features and ACT values for HI participants. From left to right: Roving condition, ACC feature, explained variance (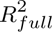), number of data points (N), and p values.

| Condition | ACC feature | $R_{full}^2$ | N | p |
| --- | --- | --- | --- | --- |
| No roving | 0 dB CL F | 0.11 | 28 | 0.08 |
| No roving | 0 dB CL Amplitude | 0.18 | 28 | 0.03 |
| No roving | 0 dB CL Latency | 0.24 | 26 | 0.01 |
| No roving | -4 dB CL F | 0.19 | 28 | 0.02 |
| No roving | -4 dB CL Amplitude | 0.11 | 28 | 0.08 |
| No roving | -4 dB CL Latency | 0.08 | 21 | 0.21 |
| No roving | -8 dB CL F | 0.17 | 28 | 0.03 |
| No roving | -8 dB CL Amplitude | 0.12 | 28 | 0.07 |
| No roving | -8 dB CL Latency | 0.36 | 17 | 0.01 |
| No roving | -12 dB CL F | 0.07 | 28 | 0.19 |
| No roving | -12 dB CL Amplitude | 0.03 | 28 | 0.39 |
| Roving | 0 dB CL F | 0.18 | 28 | 0.03 |
| Roving | 0 dB CL Amplitude | 0.21 | 28 | 0.01 |
| Roving | 0 dB CL Latency | 0.16 | 18 | 0.11 |
| Roving | -4 dB CL F | 0.13 | 28 | 0.06 |
| Roving | -4 dB CL Amplitude | 0.27 | 28 | <0.01 |
| Roving | -4 dB CL Latency | 0.38 | 14 | 0.02 |
| Roving | -8 dB CL F | 0.08 | 28 | 0.13 |
| Roving | -8 dB CL Amplitude | 0.00 | 28 | 0.96 |
| Roving | -12 dB CL F | 0.01 | 28 | 0.71 |
| Roving | -12 dB CL Amplitude | 0.00 | 28 | 0.93 |
| Δ Roving | 0 dB CL F | 0.08 | 28 | 0.14 |
| Δ Roving | 0 dB CL Amplitude | 0.03 | 28 | 0.39 |
| Δ Roving | -4 dB CL F | 0.02 | 28 | 0.44 |
| Δ Roving | -4 dB CL Amplitude | 0.05 | 28 | 0.28 |
| Δ Roving | -8 dB CL F | 0.04 | 28 | 0.3 |
| Δ Roving | -8 dB CL Amplitude | 0.13 | 28 | 0.06 |
| Δ Roving | -12 dB CL F | 0.05 | 28 | 0.27 |
| Δ Roving | -12 dB CL Amplitude | 0.04 | 28 | 0.33 |
| No roving | F slope [log2/CL dB] | 0.01 | 28 | 0.59 |
| No roving | Amplitude slope [log2/CL dB] | 0.03 | 28 | 0.36 |
| Roving | F slope [log2/CL dB] | 0.20 | 28 | 0.02 |
| Roving | Amplitude slope [log2/CL dB] | 0.27 | 28 | <0.01 |
| Δ Roving | F slope [log2/CL dB] | 0.11 | 28 | 0.09 |
| Δ Roving | Amplitude slope [log2/CL dB] | 0.10 | 28 | 0.09 |

Analyses at 0 dB CL included participants from Experiments 2 and 3, whereas analyses at lower CLs and those involving slope measures were restricted to participants from Experiment 3. Across all features, the strength of the association with ACT values varied substantially. The most consistent relationships were observed for F-values and amplitudes measured at 0 dB CL. Although response latencies were also significantly associated with ACT values, latency analyses were based on fewer observations because only responses with significant detections (p < 0.01) were included. Models based on latency estimates, particularly at lower contrast levels, should therefore be interpreted cautiously because the response-detection criterion substantially reduced the number of available observations.

For recordings obtained at 0 dB CL, both F-values and amplitudes showed significant associations with ACT values (Figure 8). Importantly, these relationships were strengthened when temporal roving was employed.

**Figure 8.**
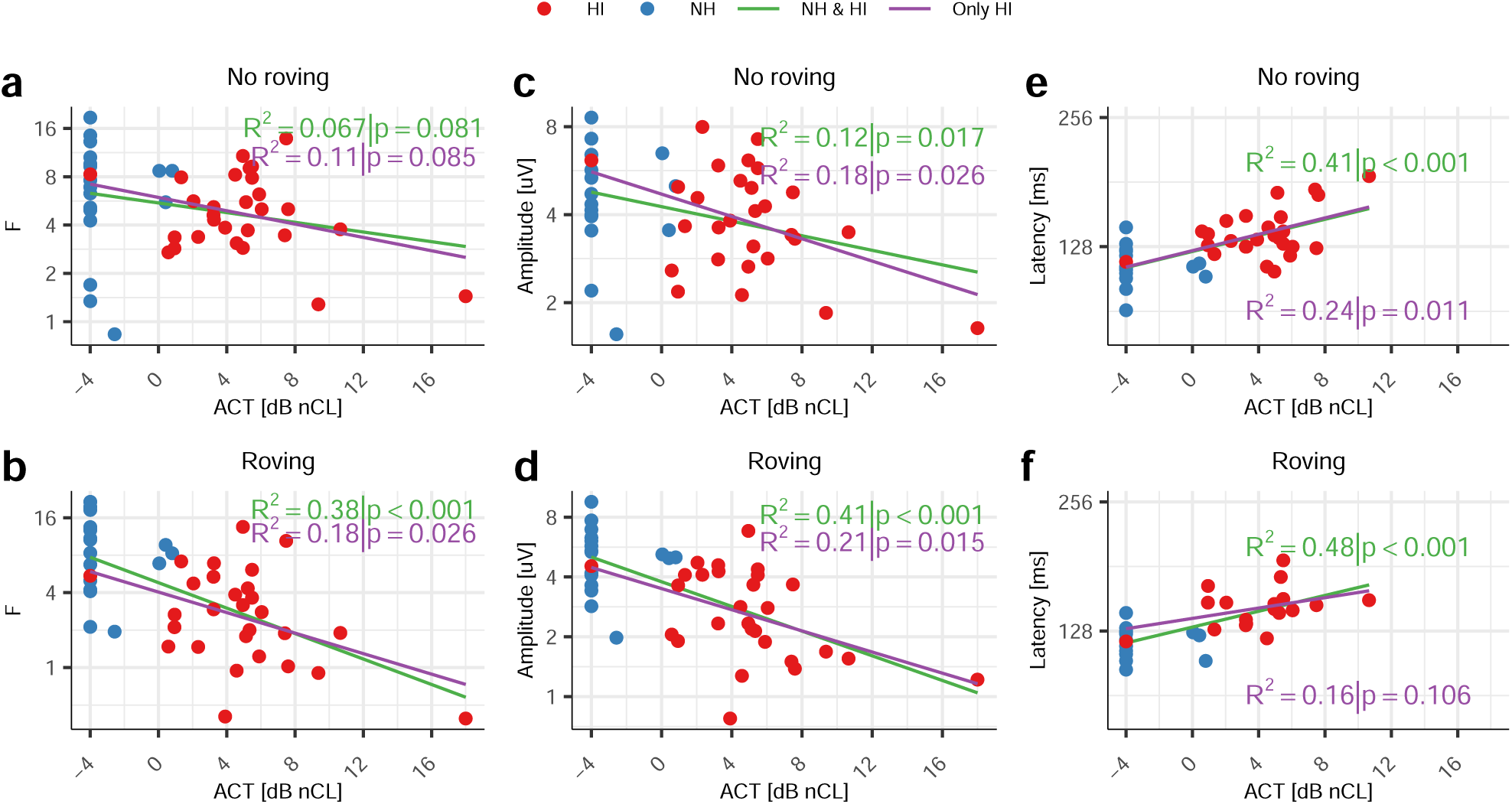
Relationship between ACT values, expressed in dB normalised contrast level (dB nCL), and ACC response strength (**a**, no roving; **b**, roving), N1-P2 amplitude (**c**, no roving; **d**, roving), and N1 latency (**e**, no roving; **f**, roving; only responses with p < 0.01 were included). Each symbol represents an individual participant. NH listeners are shown in blue and HI listeners in red. Separate regressions were fitted for the combined NH + HI dataset (green line) and for the HI group alone (purple line).

For F-values, the proportion of explained variance increased from approximately 7% in the no-roving condition to 38% in the roving condition (NH + HI). Similarly, explained variance for amplitudes increased from approximately 12% without roving to 41% with roving. Significant associations were also observed when analyses were restricted to HI participants, although the proportion of explained variance was generally reduced.

Slope measures exhibited a similar pattern. Significant relationships with ACT values were observed for F-value slopes (NH + HI: *R*^2^ = 0.25, p = 0.001; HI only: *R*^2^ = 0.20, p = 0.018) and amplitude slopes (NH + HI: *R*^2^ = 0.23, p = 0.002; HI only: *R*^2^ = 0.27, p = 0.005), but only when derived from responses obtained under temporal roving conditions (Sup. Figure 4). In contrast, features based on differences between roving and no-roving responses were generally poor predictors of behavioural performance, likely reflecting the increased measurement variability associated with difference scores.

Having established a significant relationship between cortical responses and behavioural thresholds, we next examined whether predictive performance could be improved by incorporating hearing sensitivity and participant age. Models were therefore fitted using ACC features together with PTA from the better ear and age as predictors. Full models initially included all predictors and their interactions before model reduction (see Methods). The resulting analyses are summarised in Table 3 and Table 4.

**Table 3.** Final linear models after stepwise reductions for NH and HI participants. From left to right: Roving condition, ACC feature, explained variance for the reduced (*R*^2^) and full (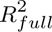) models, number of data points (N), and final model coefficients. Only model terms whose removal resulted in a significant reduction in explained variance (F-test) were retained.

| Condition | ACC feature | $R^2$ | $R^2_{full}$ | N | Intercept | PTA | Age | ACC | Age:PTA | ACC:PTA | ACC:Age | ACC:PTA:Age |
| --- | --- | --- | --- | --- | --- | --- | --- | --- | --- | --- | --- | --- |
| No roving | 0 dB CL F | 0.80 | 0.81 | 46 | -0.87 | 0.37 | -0.11 | -1.69 | N.S. | -0.09 | 0.07 | N.S. |
| No roving | 0 dB CL Amplitude | 0.75 | 0.79 | 46 | -4.42 | 0.29 | N.S. | 0.92 | N.S. | -0.04 | N.S. | N.S. |
| No roving | 0 dB CL Latency | 0.70 | 0.76 | 41 | -50.09 | N.S. | 0.12 | 6.4 | N.S. | N.S. | N.S. | N.S. |
| No roving | -4 dB CL F | 0.79 | 0.80 | 38 | 1.08 | 0.28 | -0.09 | -2.89 | N.S. | -0.08 | 0.08 | N.S. |
| No roving | -4 dB CL Amplitude | 0.70 | 0.76 | 38 | -2.78 | 0.24 | N.S. | 0.36 | N.S. | -0.03 | N.S. | N.S. |
| No roving | -4 dB CL Latency | 0.64 | 0.70 | 29 | -7.53 | N.S. | 0.16 | N.S. | N.S. | N.S. | N.S. | N.S. |
| No roving | -8 dB CL F | 0.77 | 0.78 | 38 | -7.32 | 0.58 | 0.07 | 1.26 | 0 | -0.06 | N.S. | N.S. |
| No roving | -8 dB CL Amplitude | 0.70 | 0.75 | 38 | -3.77 | 0.26 | N.S. | 0.86 | N.S. | -0.04 | N.S. | N.S. |
| No roving | -8 dB CL Latency | 0.70 | 0.77 | 25 | -68.20 | N.S. | 0.1 | 8.94 | N.S. | N.S. | N.S. | N.S. |
| No roving | -12 dB CL F | 0.69 | 0.72 | 38 | -2.16 | 0.2 | N.S. | N.S. | N.S. | N.S. | N.S. | N.S. |
| No roving | -12 dB CL Amplitude | 0.69 | 0.73 | 38 | -2.16 | 0.2 | N.S. | N.S. | N.S. | N.S. | N.S. | N.S. |
| No roving | -12 dB CL Latency | 0.48 | 0.80 | 13 | -6.16 | N.S. | 0.13 | N.S. | N.S. | N.S. | N.S. | N.S. |
| Roving | 0 dB CL F | 0.82 | 0.83 | 46 | -0.84 | 0.24 | -0.04 | -1.94 | N.S. | -0.07 | 0.06 | N.S. |
| Roving | 0 dB CL Amplitude | 0.82 | 0.83 | 46 | 2.78 | 0.36 | -0.14 | -4.15 | N.S. | -0.16 | 0.12 | N.S. |
| Roving | 0 dB CL Latency | 0.64 | 0.74 | 34 | -7.00 | N.S. | 0.15 | N.S. | N.S. | N.S. | N.S. | N.S. |
| Roving | -4 dB CL F | 0.78 | 0.80 | 38 | -0.82 | 0.29 | -0.08 | -2.34 | N.S. | -0.1 | 0.08 | N.S. |
| Roving | -4 dB CL Amplitude | 0.77 | 0.78 | 38 | -8.69 | 0.66 | 0.07 | 2.16 | -0.01 | -0.09 | N.S. | N.S. |
| Roving | -4 dB CL Latency | 0.68 | 0.80 | 21 | -146.63 | N.S. | N.S. | 20.91 | N.S. | N.S. | N.S. | N.S. |
| Roving | -8 dB CL F | 0.69 | 0.72 | 38 | -2.16 | 0.2 | N.S. | N.S. | N.S. | N.S. | N.S. | N.S. |
| Roving | -8 dB CL Amplitude | 0.69 | 0.76 | 38 | -2.16 | 0.2 | N.S. | N.S. | N.S. | N.S. | N.S. | N.S. |
| Roving | -8 dB CL Latency | 0.55 | 0.71 | 15 | -6.83 | N.S. | 0.15 | N.S. | N.S. | N.S. | N.S. | N.S. |
| Roving | -12 dB CL F | 0.69 | 0.73 | 38 | -2.16 | 0.2 | N.S. | N.S. | N.S. | N.S. | N.S. | N.S. |
| Roving | -12 dB CL Amplitude | 0.69 | 0.72 | 38 | -2.16 | 0.2 | N.S. | N.S. | N.S. | N.S. | N.S. | N.S. |
| Roving | -12 dB CL Latency | 0.63 | 0.99 | 9 | -6.00 | N.S. | 0.14 | N.S. | N.S. | N.S. | N.S. | N.S. |
| Δ Roving | 0 dB CL F | 0.56 | 0.59 | 28 | 2.75 | 0.04 | N.S. | -4.25 | N.S. | 0.13 | N.S. | N.S. |
| Δ Roving | 0 dB CL Amplitude | 0.43 | 0.49 | 28 | -0.96 | 0.17 | N.S. | N.S. | N.S. | N.S. | N.S. | N.S. |
| Δ Roving | -4 dB CL F | 0.43 | 0.54 | 28 | -0.96 | 0.17 | N.S. | N.S. | N.S. | N.S. | N.S. | N.S. |
| Δ Roving | -4 dB CL Amplitude | 0.43 | 0.54 | 28 | -0.96 | 0.17 | N.S. | N.S. | N.S. | N.S. | N.S. | N.S. |
| Δ Roving | -8 dB CL F | 0.60 | 0.65 | 28 | -1.83 | 0.2 | N.S. | 3.28 | N.S. | -0.11 | N.S. | N.S. |
| Δ Roving | -8 dB CL Amplitude | 0.59 | 0.64 | 28 | -0.60 | 0.17 | N.S. | 2.94 | N.S. | -0.11 | N.S. | N.S. |
| Δ Roving | -12 dB CL F | 0.43 | 0.57 | 28 | -0.96 | 0.17 | N.S. | N.S. | N.S. | N.S. | N.S. | N.S. |
| Δ Roving | -12 dB CL Amplitude | 0.43 | 0.50 | 28 | -0.96 | 0.17 | N.S. | N.S. | N.S. | N.S. | N.S. | N.S. |
| No roving | F-value slope | 0.83 | 0.83 | 38 | 2.44 | -0.26 | -0.15 | -62.59 | 0.01 | 5.62 | 1.73 | -0.1 |
| No roving | Amplitude slope [log2/CL dB] | 0.78 | 0.80 | 38 | -4.17 | -0.32 | 0.05 | 4.13 | 0.01 | 8.26 | N.S. | -0.12 |
| Roving | F-value slope | 0.80 | 0.82 | 38 | 1.06 | 0.25 | -0.07 | -37.45 | N.S. | -1.13 | 1 | N.S. |
| Roving | Amplitude slope [log2/CL dB] | 0.72 | 0.79 | 38 | -1.94 | 0.21 | N.S. | 1.52 | N.S. | -0.38 | N.S. | N.S. |
| Δ Roving | F slope [log2/CL dB] | 0.70 | 0.76 | 38 | -2.31 | 0.19 | N.S. | -3.32 | N.S. | 0.24 | N.S. | N.S. |
| Δ Roving | Amplitude slope [log2/CL dB] | 0.69 | 0.73 | 38 | -2.16 | 0.2 | N.S. | N.S. | N.S. | N.S. | N.S. | N.S. |

**Table 4.** Final linear models after stepwise reductions in HI participants. From left to right: Roving condition, ACC feature, explained variance for the reduced (*R*^2^) and full (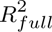) models, number of data points (N), and final model coefficients. Only model terms whose removal resulted in a significant reduction in explained variance (F-test) were retained.

| Condition | ACC feature | $R^2$ | $R_{full}^2$ | N | Intercept | Age | PTA | ACC | Age:PTA | ACC:Age | ACC:PTA | ACC:PTA:Age |
| --- | --- | --- | --- | --- | --- | --- | --- | --- | --- | --- | --- | --- |
| No roving | 0 dB CL F | 0.79 | 0.79 | 28 | 177.16 | -2.59 | -2.63 | -84.69 | 0.04 | 1.23 | 1.54 | -0.02 |
| No roving | 0 dB CL Amplitude | 0.55 | 0.71 | 28 | -6.48 | N.S. | 0.35 | 3.17 | N.S. | N.S. | -0.11 | N.S. |
| No roving | 0 dB CL Latency | 0.17 | 0.46 | 26 | 1.10 | N.S. | 0.1 | N.S. | N.S. | N.S. | N.S. | N.S. |
| No roving | -4 dB CL F | 0.67 | 0.74 | 28 | -5.83 | N.S. | 0.29 | 3.18 | N.S. | N.S. | -0.09 | N.S. |
| No roving | -4 dB CL Amplitude | 0.69 | 0.69 | 28 | 269.77 | -3.82 | -5.67 | -142.79 | 0.08 | 2.03 | 3.21 | -0.05 |
| No roving | -4 dB CL Latency | 0.56 | 0.56 | 21 | -3731.22 | 54.75 | 68.77 | 519.08 | -1.03 | -7.61 | -9.55 | 0.14 |
| No roving | -8 dB CL F | 0.77 | 0.77 | 28 | 93.66 | -1.34 | -1.45 | -76.53 | 0.02 | 1.11 | 1.56 | -0.02 |
| No roving | -8 dB CL Amplitude | 0.54 | 0.67 | 28 | -7.49 | N.S. | 0.38 | 4.78 | N.S. | N.S. | -0.17 | N.S. |
| No roving | -8 dB CL Latency | 0.36 | 0.63 | 17 | -91.11 | N.S. | N.S. | 13.08 | N.S. | N.S. | N.S. | N.S. |
| No roving | -12 dB CL F | 0.63 | 0.63 | 28 | 60.87 | -0.9 | -0.74 | -95.86 | 0.01 | 1.39 | 2.21 | -0.03 |
| No roving | -12 dB CL Amplitude | 0.43 | 0.54 | 28 | -0.96 | N.S. | 0.17 | N.S. | N.S. | N.S. | N.S. | N.S. |
| Roving | 0 dB CL F | 0.66 | 0.76 | 28 | -3.20 | N.S. | 0.23 | 2.4 | N.S. | N.S. | -0.08 | N.S. |
| Roving | 0 dB CL Amplitude | 0.75 | 0.75 | 28 | 210.79 | -2.99 | -3.87 | -114.38 | 0.06 | 1.64 | 2.19 | -0.03 |
| Roving | 0 dB CL Latency | 0.00 | 0.48 | 18 | 3.95 | N.S. | N.S. | N.S. | N.S. | N.S. | N.S. | N.S. |
| Roving | -4 dB CL F | 0.83 | 0.83 | 28 | 78.99 | -1.17 | -1.18 | -75.91 | 0.02 | 1.11 | 1.94 | -0.03 |
| Roving | -4 dB CL Amplitude | 0.80 | 0.80 | 28 | 258.02 | -3.59 | -5.35 | -158.31 | 0.08 | 2.23 | 3.55 | -0.05 |
| Roving | -4 dB CL Latency | 0.51 | 0.82 | 14 | -109.53 | 1.6 | 4.44 | N.S. | -0.06 | N.S. | N.S. | N.S. |
| Roving | -8 dB CL F | 0.72 | 0.72 | 28 | 32.47 | -0.55 | -0.17 | -99.12 | 0.01 | 1.48 | 2.13 | -0.03 |
| Roving | -8 dB CL Amplitude | 0.60 | 0.63 | 28 | 32.47 | -0.49 | 0.19 | -32.04 | N.S. | 0.46 | N.S. | N.S. |
| Roving | -12 dB CL F | 0.55 | 0.63 | 28 | -1.81 | N.S. | 0.19 | 7.17 | N.S. | N.S. | -0.21 | N.S. |
| Roving | -12 dB CL Amplitude | 0.43 | 0.53 | 28 | -0.96 | N.S. | 0.17 | N.S. | N.S. | N.S. | N.S. | N.S. |
| Δ Roving | 0 dB CL F | 0.56 | 0.59 | 28 | 2.75 | N.S. | 0.04 | -4.25 | N.S. | N.S. | 0.13 | N.S. |
| Δ Roving | 0 dB CL Amplitude | 0.43 | 0.49 | 28 | -0.96 | N.S. | 0.17 | N.S. | N.S. | N.S. | N.S. | N.S. |
| Δ Roving | -4 dB CL F | 0.43 | 0.54 | 28 | -0.96 | N.S. | 0.17 | N.S. | N.S. | N.S. | N.S. | N.S. |
| Δ Roving | -4 dB CL Amplitude | 0.43 | 0.54 | 28 | -0.96 | N.S. | 0.17 | N.S. | N.S. | N.S. | N.S. | N.S. |
| Δ Roving | -8 dB CL F | 0.60 | 0.65 | 28 | -1.83 | N.S. | 0.2 | 3.28 | N.S. | N.S. | -0.11 | N.S. |
| Δ Roving | -8 dB CL Amplitude | 0.59 | 0.64 | 28 | -0.60 | N.S. | 0.17 | 2.94 | N.S. | N.S. | -0.11 | N.S. |
| Δ Roving | -12 dB CL F | 0.43 | 0.57 | 28 | -0.96 | N.S. | 0.17 | N.S. | N.S. | N.S. | N.S. | N.S. |
| Δ Roving | -12 dB CL Amplitude | 0.43 | 0.50 | 28 | -0.96 | N.S. | 0.17 | N.S. | N.S. | N.S. | N.S. | N.S. |
| No roving | F slope [log2/CL dB] | 0.68 | 0.71 | 28 | 4.21 | -0.22 | 0.4 | 77.72 | N.S. | N.S. | -1.77 | N.S. |
| No roving | Amplitude slope [log2/CL dB] | 0.43 | 0.63 | 28 | -0.96 | N.S. | 0.17 | N.S. | N.S. | N.S. | N.S. | N.S. |
| Roving | F slope [log2/CL dB] | 0.76 | 0.76 | 28 | 110.95 | -1.56 | -1.94 | -782.2 | 0.03 | 11.19 | 16.98 | -0.25 |
| Roving | Amplitude slope [log2/CL dB] | 0.60 | 0.67 | 28 | -1.10 | N.S. | 0.17 | 30.63 | N.S. | N.S. | -1.13 | N.S. |
| Δ Roving | F slope [log2/CL dB] | 0.75 | 0.75 | 28 | 34.19 | -0.45 | -0.06 | 1418.23 | 0 | -20.23 | -35.03 | 0.5 |
| Δ Roving | Amplitude slope [log2/CL dB] | 0.43 | 0.56 | 28 | -0.96 | N.S. | 0.17 | N.S. | N.S. | N.S. | N.S. | N.S. |

When considering all participants (NH and HI), PTA and age accounted for a substantial proportion of the variance in ACT values, explaining approximately 73% and 60%, respectively. Nonetheless, the inclusion of ACC measures consistently improved model performance. For ACC features measured at 0 dB CL (n = 46), full models explained between 82% and 83% of the variance under roving conditions and between 79% and 81% under no-roving conditions (Figure 9a&b). For most models based on ACC features measured at 0 dB CL, the ACC feature was retained during stepwise model reduction because its removal produced a significant reduction in explained variance. These findings indicate that ACC-derived measures generally contributed information beyond that accounted for by PTA, age, and the retained interaction terms (Table 3).

**Figure 9.**
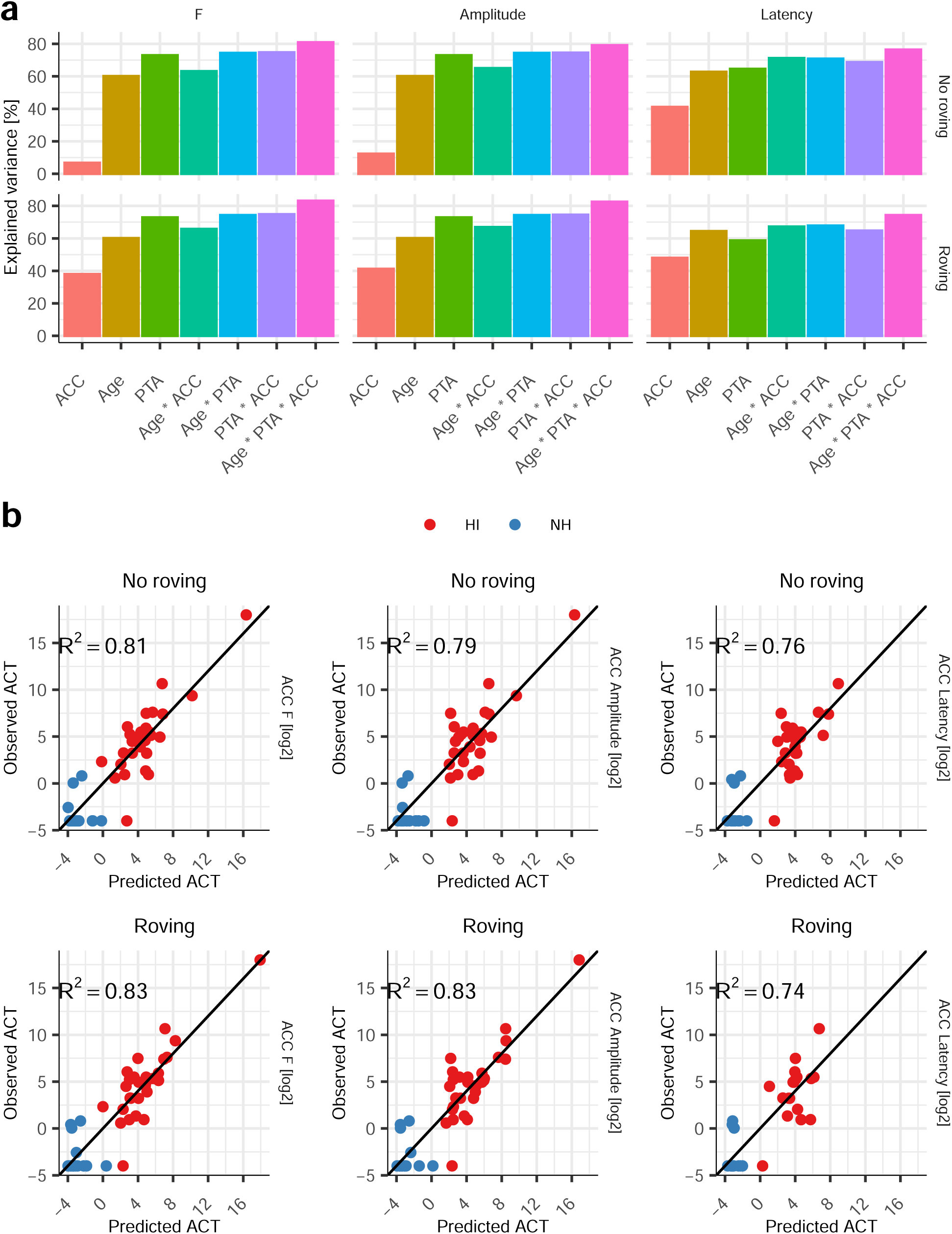
Prediction of ACT values for NH and HI listeners. **a**, Variance explained by a series of regression models incorporating ACC features measured at 0 dB CL (strength—F-value—, amplitude, or latency), age, better ear PTA, and their interactions. ACC features were obtained without temporal roving (top) or with temporal roving (bottom). **b**, Predicted versus observed ACT values for the models incorporating all predictors. Participant groups are colour coded (NH, blue; HI, red).

Similar improvements were observed for models incorporating slope measures. For example, models incorporating F-value slopes alongside PTA and age (n = 38) explained between 72% and 83% of the variance, compared with 69% and 49% explained by PTA and age alone, respectively. Features derived from differences between roving and no-roving conditions again yielded less consistent results (Table 3).

Because NH participants were generally younger and exhibited better hearing sensitivity than HI participants, analyses were repeated using only the HI cohort (n = 28). Within this group, PTA remained a significant predictor, accounting for approximately 43% of the variance in ACT values, whereas age contributed relatively little additional explanatory power (approximately 2%).

Importantly, several ACC features continued to provide substantial predictive value. Models incorporating ACC measures at 0 dB CL explained between 46% (latency under roving) and 79% (F-value under no-roving conditions) of the variance in ACT values (Figure 10). Likewise, models based on ACC slopes explained between 43% (amplitude slope under no-roving conditions) and 76% (F-value slope under roving conditions) of the variance. Across analyses, F-value-derived metrics generally provided the strongest and most consistent associations with behavioural performance.

**Figure 10.**
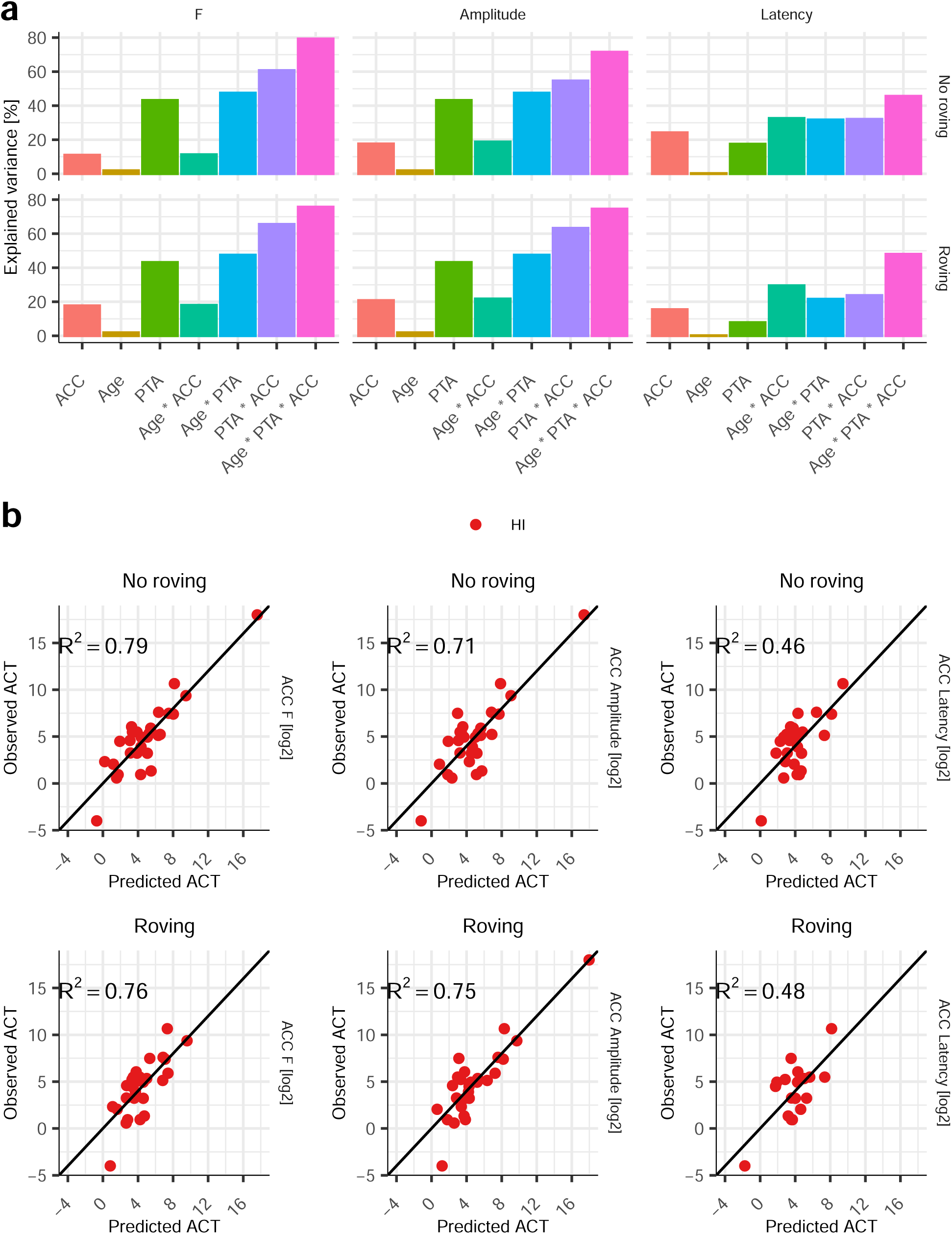
Prediction of ACT performance from ACC measures in HI listeners. **a**, Variance explained by a series of regression models incorporating ACC features measured at 0 dB CL (strength—F-value—, amplitude, or latency), age, better ear PTA, and their interactions. ACC features were obtained without temporal roving (top) or with temporal roving (bottom). **b**, Predicted versus observed ACT values for the model incorporating all predictors. Only HI listeners were included in this analysis.

Taken together, these findings demonstrate a robust relationship between cortical discrimination responses and behavioural discrimination thresholds. Although hearing sensitivity, as quantified by PTA, explained a substantial proportion of the variance in ACT values, several ACC-derived measures accounted for additional variance beyond that explained by audiometric thresholds and age alone. The strongest associations were observed for response-strength measures obtained under temporal-roving conditions, suggesting that these measures provide particularly sensitive indicators of perceptual discrimination ability.

## Discussion

### 4.1. Acoustic context strongly influences cortical discrimination responses

The present study investigated how acoustic context, temporal-envelope variability, and hearing loss influence cortical representations of spectral discrimination. Across experiments, ACC responses were strongly modulated by the statistical properties of the stimulus, indicating that cortical discrimination responses cannot be understood solely in terms of the physical magnitude of an acoustic change. Rather, the neural representation of spectral contrast depended on both the structure of the ongoing stimulus and the perceptual cues available to listeners.

The first experiment replicated and extended previous observations showing that ACC responses depend on carrier statistics and transition direction. Consistent with Simonsen et al. (2025), responses elicited using a TC were substantially larger than those elicited by a NC, and conventional STM stimuli produced pronounced asymmetries between reference-to-target and target-to-reference transitions. Importantly, these asymmetries were largely reduced when DSTMs were employed, producing more symmetrical responses across transition directions. Together, these findings indicate that cortical sensitivity to a spectral change is strongly influenced by the spectrotemporal context in which that change occurs.

An additional factor likely contributing to the larger responses observed for DSTM stimuli is the abrupt nature of the spectral transitions. Unlike conventional STM stimuli, for which spectral and temporal structure evolve continuously over time, DSTM stimuli contain discrete changes in spectral configuration. Similar abrupt spectral transitions have previously been shown to evoke robust ACC responses and to be associated with behavioural speech perception (Sohier et al., 2021). The present findings therefore suggest that cortical discrimination mechanisms may be particularly sensitive to sudden changes in spectral structure, consistent with the role of auditory cortex in detecting behaviourally relevant acoustic events. The observed effects are broadly consistent with context-dependent coding mechanisms. Auditory neurons continuously adjust their responsiveness according to the statistical properties of recent sensory inputs, including both spectral variance and stimulus-level distributions (Dean et al., 2005; Kvale and Schreiner, 2004; Nagel and Doupe, 2006; Dean et al., 2008; Wen et al., 2009; Wen et al., 2012; Herrmann et al., 2013a; Herrmann et al., 2013b; Herrmann et al., 2014; Bakay et al., 2018; Espejo et al., 2019; Willmore and King, 2022). Such adaptive processes have been observed throughout the auditory pathway and are thought to optimise neural sensitivity within the prevailing sensory environment. Under this framework, the reduced responses observed for conventional STM stimuli may reflect the greater spectrotemporal variability present within the target segment, which alters the neural representation against which subsequent acoustic changes are evaluated. The same mechanism may also account for the smaller responses observed in the NC condition relative to the TC condition. Specifically, the intrinsic envelope fluctuations and associated spectrotemporal variability of the NC stimulus may increase variability in the sensory input, thereby broadening or weakening the neural representation of the ongoing context and reducing the salience of subsequent changes.

Although increasing the alternation interval increased overall ACC strength, the fundamental effects of carrier type and stimulus structure remained unchanged. This finding argues against a simple explanation based solely on response-rate adaptation or neural refractoriness. Instead, the persistence of these effects across stimulation rates suggests that acoustic context exerts a stronger influence on cortical discrimination responses than the temporal spacing between successive transitions. Collectively, these results demonstrate that acoustic context is a major determinant of cortical sensitivity to spectrotemporal change.

### 4.2. Temporal-envelope variability dissociates intensity and spectral contributions to cortical discrimination

One of the most striking findings of the present study emerged from Experiment 2. Introducing speech-like temporal-envelope variability strongly attenuated ACC responses elicited by intensity changes whilst largely preserving, and in some cases slightly enhancing, responses elicited by spectral changes. This dissociation suggests that temporal-envelope variability influences the cortical processing of intensity and spectral information in fundamentally different ways.

In the absence of spectral cues, intensity transitions elicited robust ACCs, confirming that relatively small changes in sound level are sufficient to produce pronounced cortical discrimination responses. This observation is consistent with previous ACC studies demonstrating strong sensitivity to intensity change (Martin and Boothroyd, 2000; Dimitrijevic et al., 2011; Nishihara et al., 2011; Fujita et al., 2022). Importantly, however, introducing temporal roving markedly reduced the detectability of intensity-driven responses whilst leaving spectral-cue responses largely intact. Because the spectral manipulation itself remained unchanged across roving conditions, the observed effects are unlikely to reflect differences in stimulus salience and instead point toward changes in the contextual representation against which discrimination occurs. Although the present data do not directly identify the underlying mechanism, one possible explanation is that temporal-envelope variability disrupts the formation of a stable loudness reference. Under stationary listening conditions, repeated exposure to a relatively constant acoustic context may facilitate the detection of abrupt intensity changes. In contrast, ongoing envelope fluctuations continuously alter instantaneous sound level, reducing the reliability of intensity-based contextual representations. Spectral information, however, remains largely preserved because the underlying spectral structure of the stimulus is unchanged despite the varying envelope.

The latency findings support this interpretation. Responses elicited by spectral cues exhibited shorter and more consistent latencies than responses elicited exclusively by intensity changes. Furthermore, spectral-cue responses remained relatively insensitive to temporal roving, whereas intensity-driven responses became substantially more variable. Together, these results suggest that cortical discrimination relies on partially distinct processes depending on the cue being discriminated. Importantly, the temporal-roving manipulation was designed to approximate low-frequency envelope fluctuations commonly found in speech and other natural sounds. The present findings therefore suggest that cortical discrimination mechanisms may contribute substantially to neural responses elicited by complex sounds in which supra-threshold envelope and frequency cues occur simultaneously, as is commonly the case in natural speech.

More broadly, they indicate that spectrally driven cortical responses can be partially dissociated from competing loudness-related contributions by manipulating temporal-envelope statistics.

### 4.3. Cortical representations of spectral contrast differ between normal-hearing and hearing-impaired listeners

Having established that temporal roving reduced intensity-related contributions to the ACC, Experiment 3 examined cortical sensitivity to spectral contrast in NH and HI listeners. Across both groups, ACC response strength decreased systematically as spectral contrast was reduced, demonstrating that ACC measures are sensitive to the perceptual salience of spectrotemporal differences.

However, the shape of this relationship differed markedly between listener groups. NH listeners exhibited a non-monotonic contrast-response function under stationary conditions, whereas temporal roving revealed a more orderly monotonic decrease in response strength with decreasing contrast. In contrast, HI listeners showed monotonic reductions across contrast levels regardless of roving condition and exhibited substantially weaker responses overall, particularly under roving conditions.

One possible explanation is that temporal roving alters the relative contribution of spectral and non-spectral cues to cortical change detection. Because hearing impairment is associated with reduced spectral resolution and poorer spectrotemporal sensitivity (Bernstein et al., 2013; Mehraei et al., 2014; Zaar et al., 2023; Sohier et al., 2021), HI listeners may rely more heavily on auxiliary cues arising from stimulus temporal envelope and level, spectral weighting, or across-frequency excitation patterns. This idea is supported by studies indicating that sensitivity to slow temporal-envelope fluctuations can remain relatively preserved following hearing loss and may even be enhanced under some conditions (Goossens et al., 2019; Fuglsang et al., 2020; McClaskey, 2024).

Under this interpretation, temporal roving acts as a probe of the cues contributing to cortical discrimination. Listeners with robust spectral representations should remain relatively insensitive to the removal of intensity-based contextual cues, whilst listeners relying more heavily on non-spectral information would be expected to show larger reductions in response strength. The present findings closely match this prediction.

Consistent with this interpretation, NH listeners appear capable of maintaining robust spectral representations when intensity-related contextual cues are degraded. Under these conditions, reducing non-spectral contributions may increase the relative weighting of spectral information, resulting in more monotonic contrast-response functions and, at high contrast levels, modest improvements in cortical response strength.

Interestingly, NH listeners also exhibited a larger dynamic range of cortical responses across contrast levels than HI listeners. Whilst NH responses changed substantially as spectral contrast was reduced, HI responses occupied a narrower response range and exhibited a more compressed contrast-response function. This pattern suggests that hearing loss may compress cortical representations of spectral contrast, reducing the ability of the auditory system to differentiate among varying degrees of spectral salience. Such compression could contribute to the perceptual difficulties experienced by HI listeners in complex auditory environments, where accurate representation of subtle spectral differences is essential for sound segregation and speech understanding.

The latency findings provide additional support for this interpretation. Response latencies increased systematically as spectral contrast decreased and were consistently prolonged in HI listeners, even when neither response strength nor amplitude was statistically different between groups. Longer latencies are often interpreted as reflecting reduced efficiency or reduced certainty in cortical discrimination processes. The present findings therefore suggest that hearing impairment affects not only the strength of cortical discrimination responses but also the timing with which spectrotemporal information is processed.

An alternative explanation for the reduced responses observed in HI listeners is reduced effective loudness rather than impaired spectrotemporal processing. Although audibility was ensured for HI listeners, ASSR magnitudes were lower in HI than in NH listeners. This difference is compatible with lower effective sensation levels or altered envelope encoding in the HI group (Van Eeckhoutte et al., 2016).

However, the control experiment demonstrated that reducing stimulus level by 15 dB in the NH group produced only modest reductions in ACC strength, substantially smaller than the differences observed between NH and HI listeners. Together with the audibility-compensation strategy employed throughout the study, these findings suggest that the reduced cortical responses observed in the HI group cannot be explained solely by differences in sensation level and are more consistent with impaired cortical representation of spectrotemporal contrast.

Importantly, corresponding ASSRs remained largely insensitive to spectral contrast despite clear contrast-dependent effects in the ACC. This dissociation suggests that the contrast-dependent ACC effects primarily reflect change-related cortical discrimination rather than a general reduction in phase-locked encoding of the 42-Hz stimulus envelope.

### 4.4. Cortical discrimination responses predict behavioural discrimination performance

A major objective of this study was to determine whether cortical representations of spectral contrast are related to behavioural sensitivity. Across listeners, stronger ACC responses were generally associated with lower ACT values, indicating better behavioural discrimination of spectrotemporal contrast.

The strongest associations were observed for ACC measures obtained under temporal-roving conditions. Response strength measured at the highest contrast level explained a substantial proportion of the variance in ACT performance and continued to contribute predictive information even after accounting for age and hearing sensitivity. These findings indicate that cortical discrimination measures capture aspects of auditory processing that are not fully explained by conventional audiometric thresholds.

Notably, measures describing the dynamic response of the ACC across contrast levels also showed significant relationships with behavioural performance. Participants exhibiting larger changes in cortical response strength as contrast was reduced generally demonstrated superior behavioural sensitivity. This observation suggests that the cortical representation of spectral contrast is reflected not only in response magnitude at a single contrast level but also in the dynamic range with which contrast is encoded.

The observation that temporal roving strengthened several ACC-behaviour relationships is consistent with the interpretation developed throughout the preceding experiments. By reducing competing intensity-related contributions, temporal roving may increase the specificity of ACC responses to spectral discrimination. Consequently, the resulting neural responses more closely reflect the perceptual mechanisms measured behaviourally by the ACT.

Although the present study was not designed as a clinical validation study, the observed relationships between cortical and behavioural measures suggest that context-sensitive ACC paradigms may ultimately prove useful for objective assessment of spectrotemporal processing in populations for whom conventional behavioural testing is difficult.

### 4.5. Limitations and future directions

Several limitations should be considered when interpreting the present findings. First, the NH and HI groups differed not only in hearing status but also in age. Although age was included in the behavioural analyses and ACC measures explained variance beyond that accounted for by age alone, age-related changes in cortical adaptation, temporal processing, and neural response variability may have contributed to some of the observed group differences. Future studies employing age-matched NH and HI cohorts will be important to disentangle the relative contributions of peripheral hearing loss and aging to context-dependent cortical representations of spectral contrast. In addition, the multivariable regression analyses, particularly those based on latency estimates at lower contrast levels, should be considered exploratory because some models were fitted to relatively small samples and included multiple interaction terms.

Second, the present experiments employed symmetric stimulus structures in which reference and target segments had identical durations. Because previous studies have shown that cortical change responses are influenced by stimulus history and adaptation state, future work could use asymmetric STM and DSTM paradigms in which reference and target segments differ in duration. Such manipulations would provide a more direct test of how the temporal statistics of the preceding acoustic context influence cortical discrimination responses and may help further distinguish context-dependent coding mechanisms from shorter-term response-rate adaptation effects.

Third, the relationship between cortical responses and perceptual performance was evaluated using the ACT, a behavioural measure based on spectrotemporal modulation sensitivity. Although this provided a strong conceptual link between the physiological and behavioural tasks, the ecological relevance of these cortical measures could be further assessed by comparing ACC responses with speech-recognition outcomes obtained using more realistic speech-in-noise tests. Establishing whether context-sensitive ACC measures predict performance under everyday listening conditions would help clarify their functional significance for auditory perception.

Finally, although audibility was carefully controlled through individualised amplification procedures, listeners were not tested at matched sensation levels. Consequently, differences in suprathreshold audibility may have remained across participants despite ensuring that stimuli were audible. Future studies could systematically manipulate and match sensation level across NH and HI listeners to determine the extent to which cortical representations of spectral contrast depend on the effective neural coding range available to individual listeners.

### 4.6. Conclusions

The present study demonstrates that cortical representations of spectral discrimination are strongly shaped by acoustic context. Across experiments, ACC responses depended not only on the magnitude of spectral change but also on carrier statistics, transition structure, and temporal-envelope variability. Cortical discrimination responses were largest for discrete spectral transitions, highlighting the sensitivity of auditory cortex to abrupt changes in spectral structure and emphasizing the importance of contextual factors in shaping neural representations of acoustic contrast.

A central finding was that speech-like temporal-envelope variability selectively attenuated intensity-driven cortical responses whilst largely preserving responses driven by spectral contrast, revealing a partial dissociation between intensity-driven and spectrally driven discrimination processes. These results indicate that cortical responses to complex sounds reflect an interaction between sensory cues and contextual representations rather than a simple encoding of physical stimulus change. Hearing impairment was associated with weaker, delayed, and less differentiated cortical representations of spectral contrast, particularly under conditions that reduced the availability of intensity-related cues. Importantly, several cortical response measures were significantly associated with behavioural ACT performance and contributed explanatory information beyond that provided by age and audiometric thresholds alone.

Together, these findings demonstrate that cortical discrimination responses reflect dynamic and context-dependent representations of spectrotemporal contrast. More broadly, they show that manipulating acoustic context can reveal distinct contributions of spectral and intensity cues to cortical processing, providing new insight into the neural mechanisms underlying auditory discrimination.

## Acknowledgments

This study received no external funding.

## Author contributions

J.U., M.V.E., and L.S. conceptualised and designed the experiments. I.S. and M.R. performed the behavioural and neurophysiological experiments. J.U. and I.S. analysed both behavioural and neurophysiological data. J.U. wrote the manuscript with input from all authors.

## Declaration of interests

None

## Data availability

The EEG datasets generated and analysed during this study are publicly available through the DTU Data repository (DOI: 10.11583/DTU.33952357).

## Code availability

All methods described in this study are implemented in pEEGy, an open-source Python package developed for the analysis and simulation of electroencephalography data and evoked responses. The pEEGy source code is publicly available and maintained under an open-source licence. The version used for the analyses reported here is archived with a persistent digital object identifier (10.5281/zenodo.19512653), enabling reproducibility of the results. Documentation and usage examples are provided with the software. The specific source code used to generate all figures and statistical analyses will be publicly available on the date of publication.

## Supplementary figures

**Sup. Figure 1.**
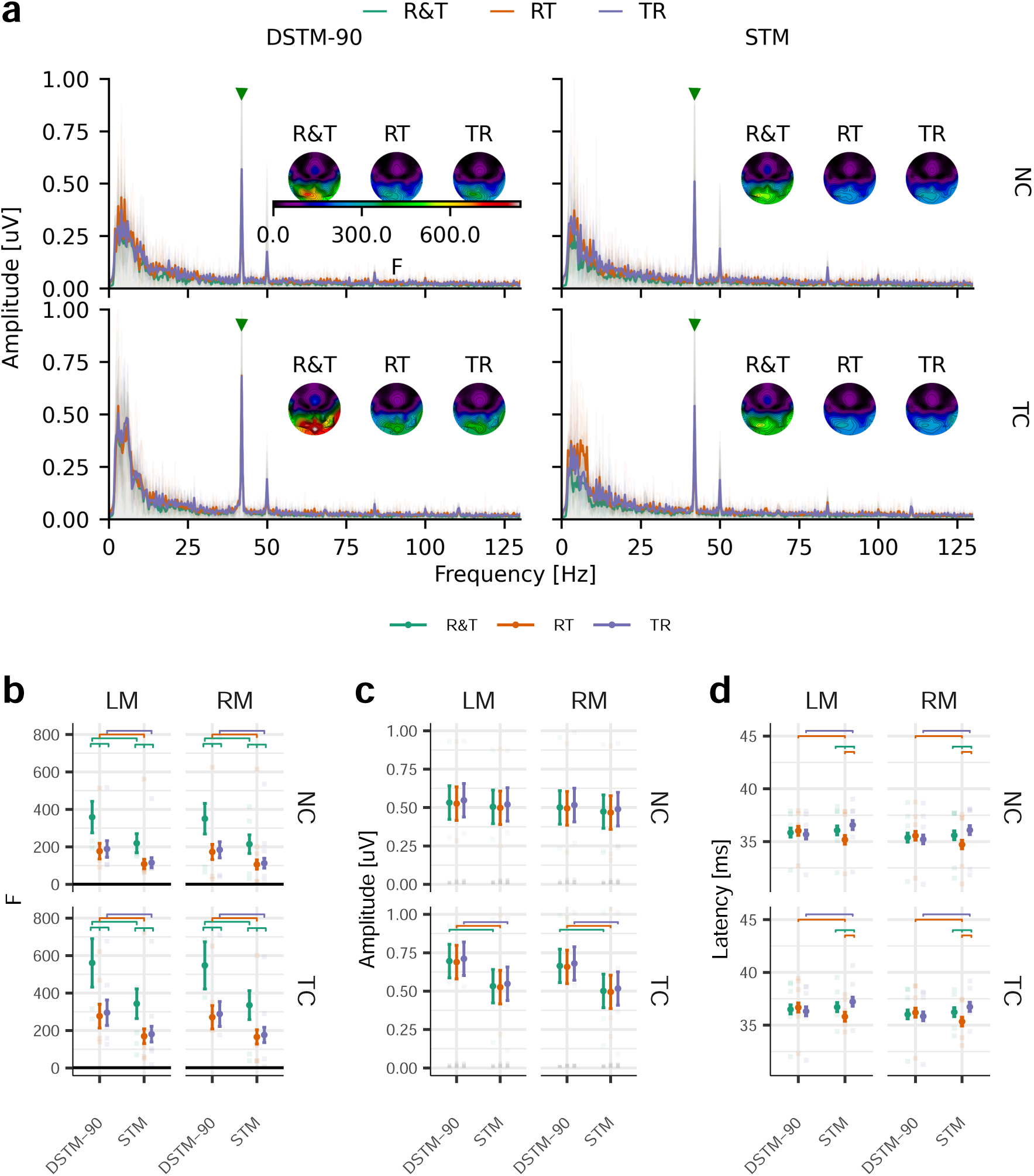
Carrier, transition direction, and channel effects on ASSRs. **a**, ASSRs evoked by DSTM-90 (left) and STM (right) stimuli presented using either a NC (top) or TC (bottom). Responses were recorded for RT, TR, and R&T transition conditions. Topographic maps illustrate ASSR response strength (F-value) across the scalp. **b**, Response strength (F-value). **c**, ASSR amplitude. **d**, ASSR latency. Individual observations are shown as semitransparent points, whilst solid symbols and error bars represent estimated marginal means ± standard error from the GLME model. Significant post hoc comparisons (FDR corrected) are indicated by horizontal bars. The horizontal black line in **b** indicates the unitary F-value. Semitransparent grey bars in **c** denote RN.

**Sup. Figure 2.**
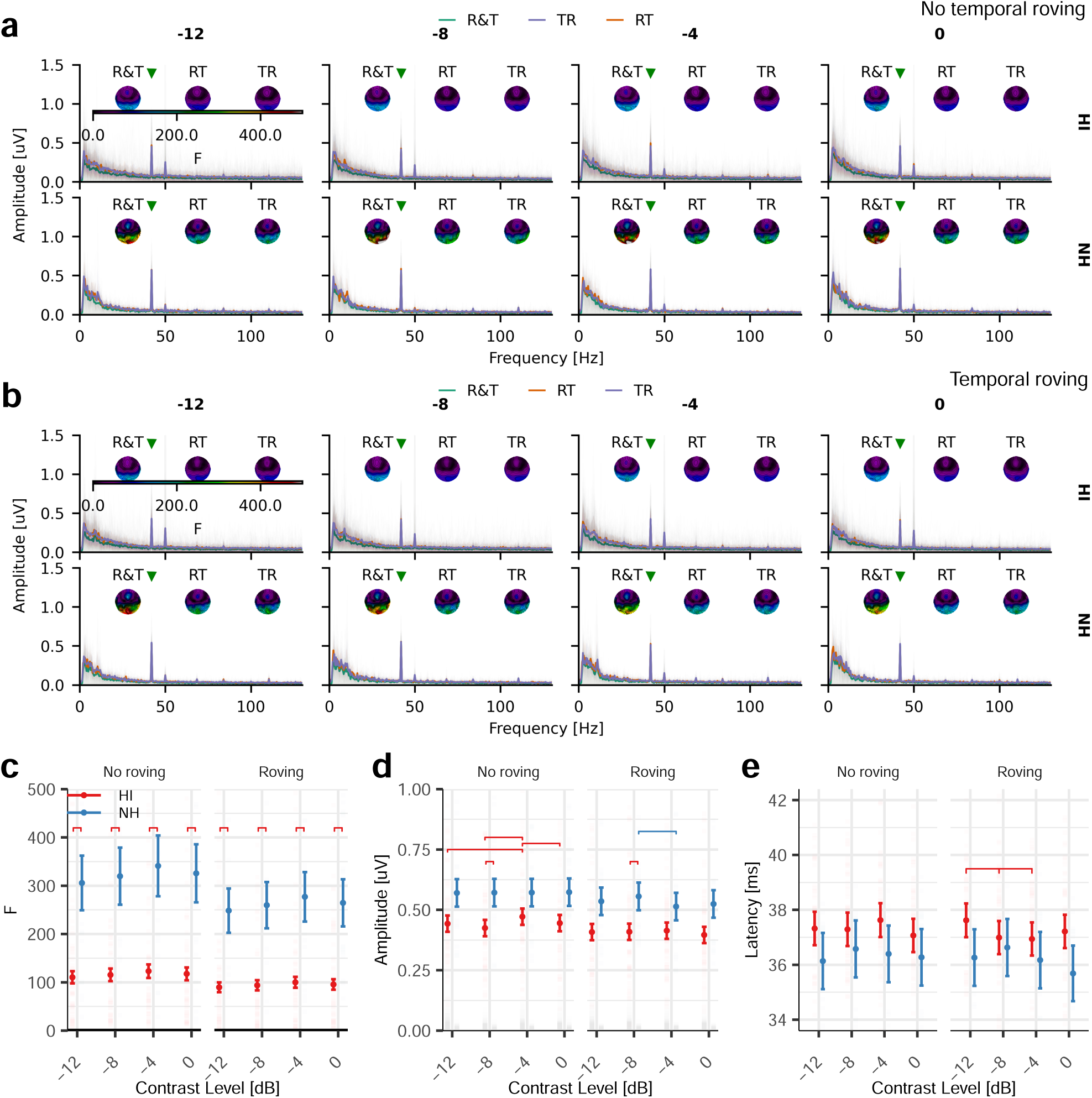
Effects of spectral contrast and temporal roving on ASSRs in NH and HI listeners. **a**, ASSR responses evoked by DSTM-90 stimuli using a TC without temporal roving. RT, TR, and combined R&T transitions are indicated by the corresponding colour. Responses are shown for HI listeners (top) and NH listeners (bottom). The spectral CL is indicated above each panel. Individual responses are shown as semitransparent traces (left and right mastoids) and grand-average responses by solid traces. Topographic maps show the average response strength (F-value) across the scalp. Vertical dashed lines indicate the analysis window. **b**, Same as **a**, but with temporal roving applied. **c**, Response strength, **d** ASSR amplitude, and **e** ASSR latency. Data in **c**-**e** represent R&T responses averaged across left and right mastoid electrodes. Individual observations are shown as semitransparent points, whilst solid symbols and error bars represent estimated marginal means ± standard error from the GLME. Significant post hoc comparisons (FDR corrected) are indicated by horizontal bars. The horizontal solid black line in **c** indicates the unitary F-value. Semitransparent grey bars in **d** denote RN.

**Sup. Figure 3.**
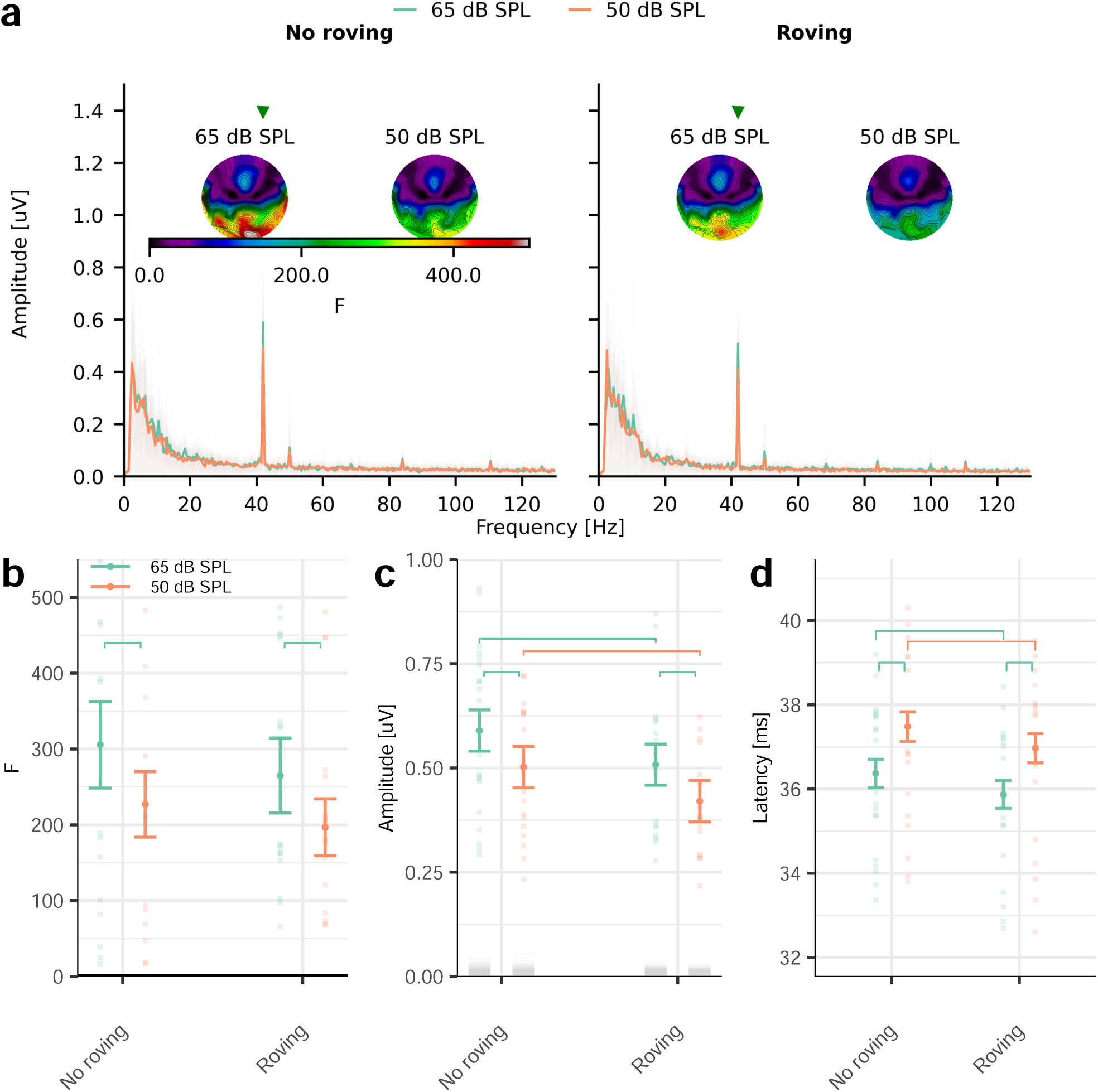
Effect of presentation level and temporal roving on ASSRs in NH listeners. **a**, ASSR waveforms evoked by DSTM-90 using a TC at presentation levels of 50 dB SPL (orange) and 65 dB SPL (green), without temporal roving (left) and with temporal roving (right). Spectral contrast was fixed at 0 dB CL. Individual responses are shown as semitransparent traces (left and right mastoids) and grand-average responses as solid traces. Topographic maps illustrate mean ASSR response strength (F-value) across the scalp. Vertical dashed lines indicate the analysis window. **b**, Response strength, **c** ASSR amplitude, and **d** ASSR latency. Data in **b**-**d** represent R&T responses averaged across mastoid electrodes. Individual observations are shown as semitransparent points, whilst solid symbols and error bars represent estimated marginal means ± standard error from the GLME. Significant post hoc comparisons (FDR corrected) are indicated by horizontal bars. The horizontal black line in **b** indicates the unitary F-value. Semitransparent grey bars in **c** denote RN.

**Sup. Figure 4.**
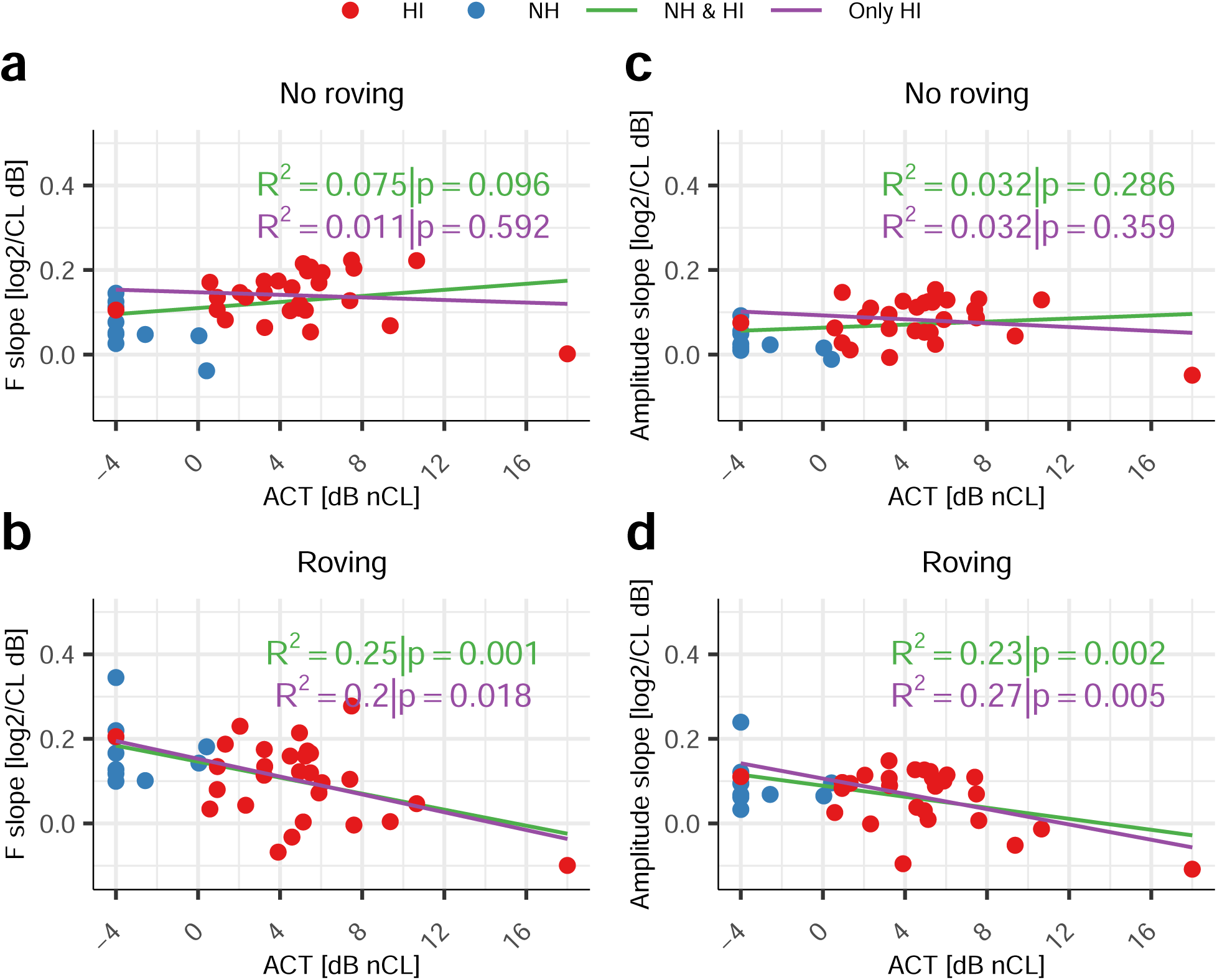
Relationship between ACT values (dB nCL) and ACC slopes for the response strength (**a** no-roving, **b** roving) and N1-P2 amplitudes (**c** no-roving, **d** roving). Each dot represents an individual participant, with dark blue dots indicating NH listeners and red dots indicating HI participants. Regressions were computed twice, including both NH and HI participants (green line), and including only HI participants (purple line).

## Supplementary tables

## Notes

### Competing Interest Statement

The authors have declared no competing interest.

